# Single-molecule imaging and tracking on clinical liquid biopsies reveals cancer biomarkers nanoscale organization and heterogeneity

**DOI:** 10.64898/2026.04.13.717683

**Authors:** Marrit M.E. Tholen, Roger Riera, Tijmen H.W. Hijzelaar, Hao Cao, Francesca Cortopassi, Meike E. Moers, Mitko Veta, Marjan J. Cruijsen, Daan van de Kerkhof, Volkher Scharnhorst, Lorenzo Albertazzi

**Affiliations:** Department of Biomedical Engineering, Institute for Complex Molecular Systems (ICMS), Eindhoven University of Technology, Eindhoven 5612AZ, The Netherlands; Medical Image Analysis, Department of Biomedical Engineering, Eindhoven University of Technology, 5600 MB Eindhoven, The Netherlands; Department of Hematology, Catharina Hospital, Eindhoven, The Netherlands; Clinical Laboratory, Catharina Hospital, Eindhoven, The Netherlands

## Abstract

Single-molecule imaging and tracking have revealed fundamental biological mechanisms at the molecular scale, yet their application to clinical research remains limited by technical complexity and sample preparation incompatible with patient-derived specimens. As a result, we lack information with molecular-scale resolution of clinically relevant biomarkers.

Here, we develop a workflow enabling Points Accumulation in Nanoscale Topography (PAINT) combined with single-particle tracking (SPT) on clinical liquid biopsies, allowing analysis of biomarker nanoscale organization at the single-molecule level in cancer patients. Our approach features a sample preparation tailored to liquid biopsies and requires no fixation, covalent labelling, or genetic modification, making single-molecule imaging compatible with hospital clinical workflows.

We demonstrate the method’s diversity by imaging liquid biopsies from blood, bone marrow aspirates, and pleural effusions across different cancer types. PAINT-SPT captures both the expression and mobility of clinically relevant membrane receptor biomarkers, revealing pronounced inter- and intra-patient heterogeneity at the molecular and cellular levels.

We discover that individual patients exhibit distinct molecular mobility fingerprints that reflect biomarker interaction states and correlate with clinical diagnostic readouts. Furthermore, these fingerprints distinguish healthy from cancer cells, enabling the development of a classifier that accurately identifies cancer cells based on their single-molecule behaviour.

Together, our results establish a route to investigate patient-derived clinical samples at the single-molecule level and open new opportunities to understand cancer biology and biomarker function beyond ensemble-averaged measurements.

---

Single-molecule imaging and super-resolution microscopy have transformed our ability to study biological structures beyond the diffraction limit of light. In fixed cells, single-molecule localization microscopy (SMLM) provides nanometre-scale information about biomolecule distribution while also enabling molecular counting. In particular, Points Accumulation in Nanoscale Topography (PAINT) pushed the resolution limit below 1 nm^1,2^ and allows for multiplexing up to 30 colours.^3^ Application of these methods generates unique information about the number of biomolecules and the spatial distribution^4,5^, colocalization^6–8^ and heterogeneity^3,7,9^. PAINT can be combined in live cells with single-particle tracking (SPT), allowing to obtain dynamic information such as molecular mobility and interactions. In particular, approaches like uPAINT^10^ and PAINT-SPT^11^ proved very powerful to study membrane receptors, revealing unique data about receptor oligomerization, activation and anchoring to cellular structures.^11–13,14^ Recently, molecular mobility has attracted attention as a phenotyping tool, providing a new dimension of analysis beyond static expression levels, allowing to identify distinct cell states, thanks to new AI-based classification algorithms.^11,15–18^

Despite these achievements, the vast majority of this work is performed in controlled models like cell lines, and is currently not used in a clinical setting. This is because the requirements of such complex techniques in terms of sample preparation, engineering labels and imaging conditions are often not compatible with the practical constraints of clinical samples like biopsies that need fast measurements, higher throughput to account for cell variability and do not allow for modification of the cells, like expression of fluorescent proteins or labelling tags. Recent pioneering reports showed how super-resolution microscopy can provide unique information from clinical samples, such as exosomes^19,20^, brain sections of Alzheimer’s tau aggregates^21^ and fixed cells from patients.^22,23^ Having these methods routinely used in clinical research will enhance our understanding of human diseases and guide the design of novel diagnostic and therapeutic approaches. However, this is still in its infancy and the adoption of these methods in clinical research is still challenging. Moreover, to the best of our knowledge, there is no report of single-molecule imaging and tracking in clinical samples, due to the lack of a live-cell imaging workflow tailored for the requirements of clinical specimens.

Here, we bridge the gap between live-cell single-molecule imaging and clinical research through a workflow that addresses key technical challenges of clinical imaging. Our methods allow us to perform PAINT-SPT on biopsies from cancer patients, revealing unique information about membrane biomarkers of diagnostic and therapeutic interest, such as molecule counting, distribution and mobility/interactions. This analysis revealed that each patient possesses a unique single-molecule mobility phenotype, with substantial differences between patients and wide heterogeneity at the molecular-, cellular-, and patient-level. The obtained data can be correlated with standard diagnostic assays such as immunohistochemistry (IHC)^24^, flow cytometry and gene sequencing^25^ providing complementary insights about receptor organisation and dynamics at the nanoscale^26^.

We tailored PAINT-SPT imaging for liquid biopsies, bodily fluids enriched in cancer biomolecules and cancer cells (circulating tumour cells, CTCs), which are emerging as a powerful, less-invasive alternative to solid biopsies. We test the robustness and wide applicability of our method by measuring biopsies from different sources (blood, bone marrow and pleural effusion) and different diseases (lung cancer and leukaemia). As a case study to prove the unique information that can be obtained with our method, we delved into single-molecule analysis of acute myeloid leukaemia (AML) from a cohort of patients from the Catharina Hospital in Eindhoven. For each patient, four biomarkers, chosen for their diagnostic and therapeutic potential, have been analysed, their expression and single-molecule mobility fingerprint extracted. We then investigated the differences of these fingerprints between: i) cells of the same patient (intra-patient heterogeneity); ii) cancer cells versus healthy cells; iii) different patients (inter-patient heterogeneity); iv) cancers with different mutation burden; v) different sites of biopsy and vi) patients and cell line models to verify the resemblance of in vitro models with human biology.

This imaging framework may open new avenues for biomarker discovery based on dynamic rather than static molecular features. More broadly, our methodology represents a step toward translating single-molecule imaging into clinically relevant research, offering a blueprint for nanoscale characterization of patient-derived samples.

## RESULTS

### Clinical PAINT-SPT workflow design and optimization

Figure 1 shows the imaging workflow developed in this work. This has been designed to match the requirements of single-molecule imaging with the practical issues related to liquid biopsies. The workflow covers the whole experiment, from sample preparation to data analysis and data mining efforts. It is crucial to standardize and optimize all steps to ensure a fair and reproducible comparison between patients. Figure 1A shows a schematic representation of the different steps, while in Fig. 1B, representative images of the different steps are reported.

**Figure 1:**
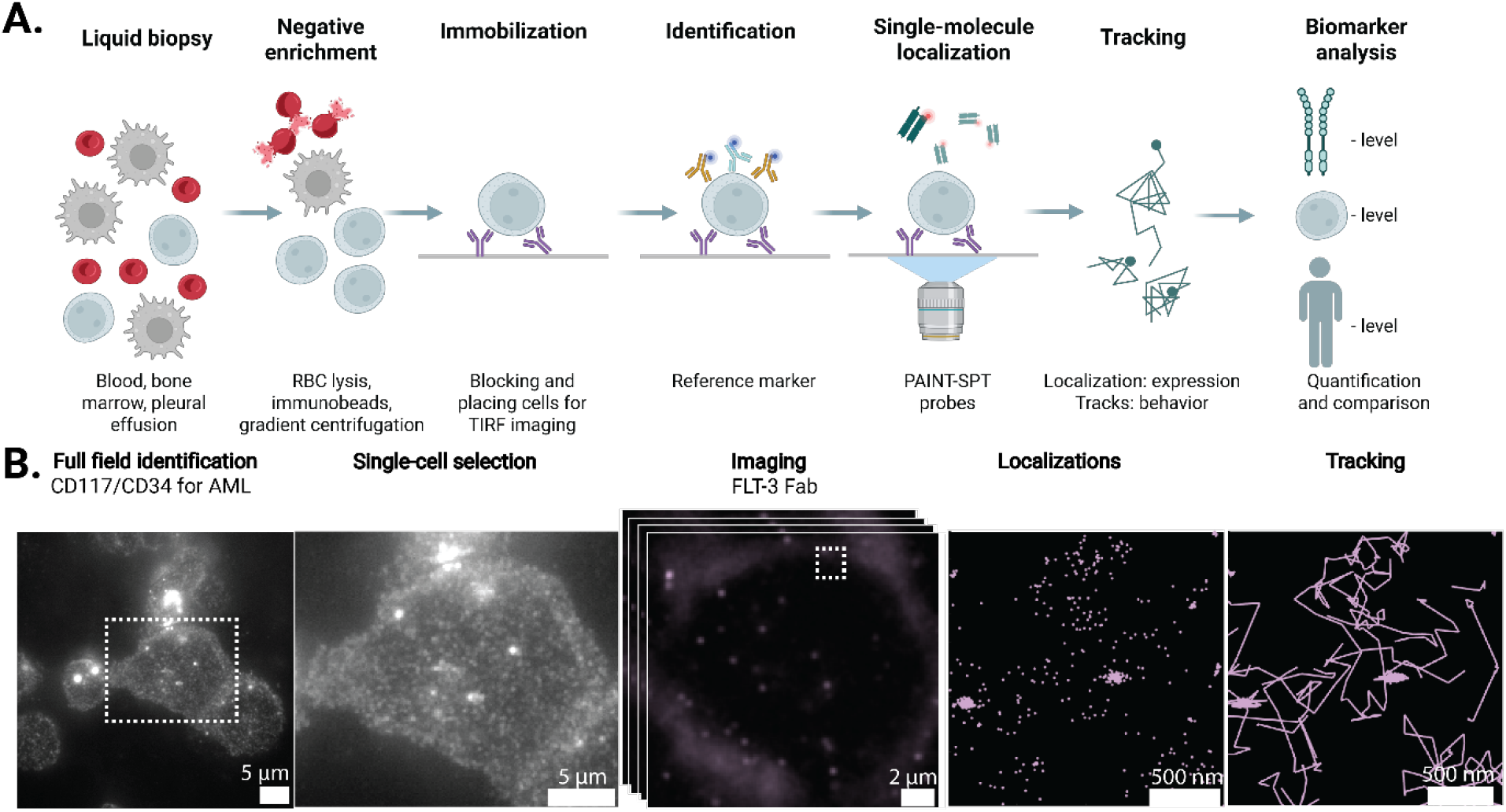
Workflow for analysis of clinical samples. A) Schematic representation of the steps in the workflow: isolation is followed by capture, identification, single-molecule imaging and single-particle tracking. B) Representative images of a cell at each step after isolation. Localisations and tracks are a zoom-in of the area indicated with the square in the events image.

### Sample preparation, cell capture and identification

A major difference of liquid biopsies compared to standard cell line samples is the presence of multiple cell types and the fact that the cells of interest (cancer cells) may be rare compared to the other cells in the mixture. ^27^ Therefore, the first challenge is to enrich the cell of interest to facilitate imaging in a later step. We explored several procedures based on positive enrichment (e.g. beads to capture and separate the cell of interest) or negative enrichment (methods to remove or lyse the undesired species), as shown in Fig. 2A-B. For bone marrow and blood samples, lysis of the red blood cells (RBCs) with ammonium chloride was sufficient to obtain a high number of desired cancer cells.^28^ For the pleural effusion from lung cancer patients, RBC removal was not sufficient as the number of cancer cells is much lower, down to a few cells per sample. Here, we compared different strategies as extensively discussed in Fig. S1,2. Overall, negative enrichment was chosen for its minimal perturbation of the sample, good viability and good recovery rate of the cancer cells as shown in Fig. 2B. Overall, these procedures proved to be robust over several types of liquid biopsies and tens of patients analysed in this study.

**Figure 2:**
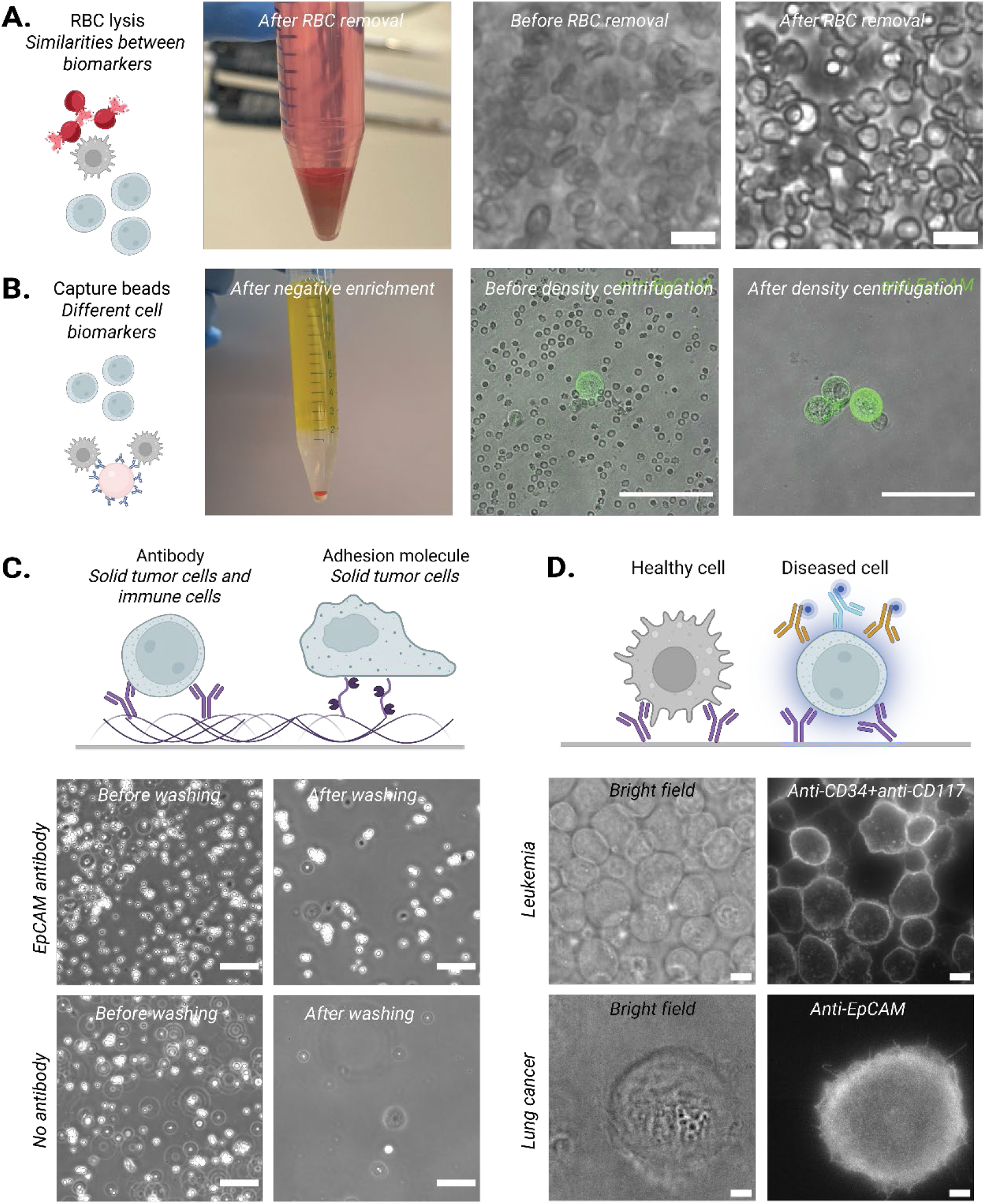
From biopsy to microscope. A) Negative enrichment is performed by RBC lysis for AML and B) density centrifugation for lung cancer, resulting in a pellet or layer with the cells of interest. In microscopy, without the RBC removal, individual cells are hard to distinguish due to the high degree of RBCs, while the use of RBC lysis buffer shows individual WBCs with a sparse RBC. Cells were washed 4 times with Leibovitz after 45 min incubation on glass. Scale bar: 20 μm. For the capture beads, the cancer cells were stained with anti-EpCAM (green). Before density centrifugation, a high number of negative cells are found around the cancer cell (WBCs, RBCs), but after density centrifugation, only the positive cells are found. The negative enrichment strategy should be chosen based on what is not damaging the cells of interest and considering patient-to-patient variability of the number of cells and marker expression. Scale bar: 100 μm C) Cells are captured on the microscopy slide using antibodies or adhesion molecules. In both the capture slide and the negative controls, the number of cells before washing is high. After washing, the cells are almost fully removed in the negative control, while they remain in the capture slide. Scale bar: 100 μm D) Identification of the malignant cells is done in the microscope. Brightfield and staining image from the same FOV, some cells were not stained for the leukaemia sample. In the pleural effusion sample, only a single positive cell fits in a field of view. Scale bar 5 μm.

To perform PAINT single-molecule imaging, the cells must be close to the glass, to enable total internal reflection (TIR) and the cells must remain immobile for the duration of the imaging. Moreover, the cell membrane should be accessible for probes. Here, we achieved this with a capture strategy based on a flow cell with a functionalized glass coverslip surface. Depending on the cell target, we optimized two protocols as shown in Fig. 2C: i) both adherent and non-adherent cells (e.g. leukaemia) can be captured on pegylated anti-fouling surfaces functionalised with antibodies against a known cancer epitope^12^, and ii) adherent cells (e.g. lung cancer) can be captured with RGD adhesion peptides without pre-knowledge of biomarkers and without bias.

In Fig. 2C can be seen that without this capturing layer, a high proportion of cells is washed away, while in the presence of the capturing layer, only a selection of cells remains after washing, indicating selectivity towards the biomarker-positive cells (Fig. 2C). Fig. S3 provides a comparison of multiple patients, captured with both anti-EpCAM and RGD surfaces, showing that RGD capture is more efficient in most cases. Overall, these capture surfaces immobilize cells at a TIR range but at the same time provide accessibility for PAINT probes for single-molecule imaging.

While enrichment and capture increase the percentage of cells of interest enormously, there is always some contamination from undesired cells that cannot be distinguished by morphology (e.g. healthy lung epithelia versus lung cancer). To solve this, we use one of the fluorescence channels (488 nm excitation) for an identification label, while super-resolution will be performed in a spectrally separated channel with minimal cross-talk (647 nm excitation). For this identification, we use the same markers used in the standard IHC diagnostic procedure (CD34 and CD117 for leukaemia, EpCAM for lung cancer), aligning the results with other clinical analyses.

In Fig. 2D, a representative image of both samples is shown, in both the brightfield and the fluorescence channel. For both samples, many morphologically identical cells can be observed in the brightfield, making identification impossible. However, when looking at the fluorescent signal, only a portion of the cells is stained, allowing for accurate identification of the cells, and allowing for the choice of where to perform single-molecule imaging. Table S1 and S2 show that the results of karyotype and immunohistochemistry in the hospital for AML and the results from our identification step matched well with clinical IHC assessments. These findings validate the effectiveness of our capture and identification workflow in clinical samples. Overall, the sample preparation reported here is robust, adaptable to virtually all types of liquid biopsies, and compatible with single-molecule imaging.

### PAINT-SPT imaging across different liquid biopsies

Common labels such as fluorescent protein or HaloTags that require genetic modification or antibodies which may alter receptor behaviour (e.g. dimerization) are not possible in clinical samples^29^ and alternative strategies such as endogenous ligands^13,15,30^ and engineered monovalent ligands^31^ are necessary. Here we tested and optimized monovalent Fab fragments, which have a lower affinity^32,33^, small size and are monovalent, ideal properties for PAINT-SPT. Fab fragments against FMS-like tyrosine kinase 3 (FLT3), T-cell immunoglobulin and mucin-domain containing-3 (TIM-3), sialic acid binding Ig-like lectin 3 (CD33), interleukin-2 receptor alpha (CD123) and programmed-death ligand-1 (PD-L1) were obtained from commercial sources or produced from validated antibodies (see materials and methods for procedure and characterization).^34,35^ All Fab domains were labelled with ATTO-643, a photostable and bright dye suited for PAINT-SPT.^13,36^ Probes were added at nanomolar concentrations in the imaging medium. The probes diffuse freely and become detectable upon binding to their target receptors on the cell surface. The probe can then be localized by Gaussian and tracked over time with millisecond time resolution.

Figure 3A shows a representative imaging result for PD-L1 imaging, an immunotherapy biomarker, in lung cancer patients. It can be appreciated how single-molecule trajectories are identified in patients positive for PD-L1 (left) while no signal is identified in PD-L1 negative samples. Overall, this shows that we can perform specific and sensitive single-molecule measurements on patients’ samples (right). The quality of the data is demonstrated by a high photon count and localization precision (Fig. 3B), comparable to standard measurements in cell lines. The resulting diffusion coefficient is also comparable to expected values.^13,37^ Overall, these results show that we can successfully perform PAINT-SPT on clinical samples with high cell specificity without compromising the quality of single-molecule localization and tracking.

**Figure 3:**
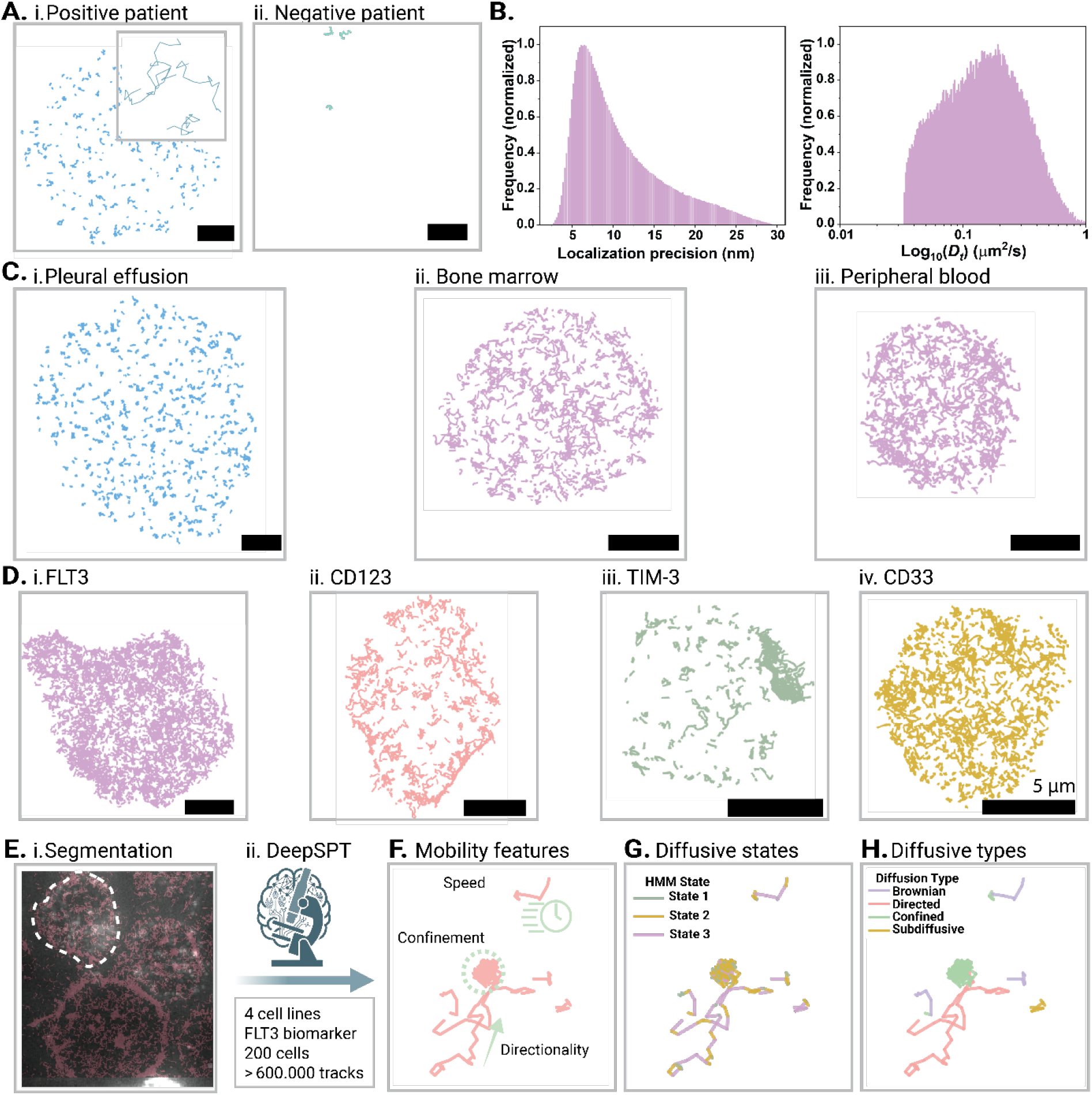
Imaging and data analysis of the workflow. A) Imaging can be performed on i) positive and ii) negative cells with high quality in B) localization precision (based on 3 random fields of view) and with biologically relevant instant diffusion coefficients. C) Imaging and tracking are performed on three different types of liquid biopsies: bone marrow, blood and pleural effusion. Scale bar 5 μm. D) Multiple different targets are imaged for AML i) FLT3, ii) CD123, iii) TIM-3 and iv) CD33. E) Data is extracted from the SPT-PAINT tracks by first i) segmenting the image, followed by ii) DeepSPT, which was trained by using 4 cell lines. This results in information on the molecular, cell and patient-levels in three ways F) mobility features, G) diffusive states and H) diffusion type prediction.

Fig. 3C shows imaging and tracking of receptors was successful for all types of liquid biopsies obtained: bone marrow (BM), peripheral blood (PB) and pleural effusion (PE). Furthermore, all five probes were successful in detecting the molecules of interest (4 for leukaemia shown in Fig. 3D and Fig. S4 and 1 for lung cancer shown in Fig. 3A). These results show the wide applicability and robustness of our workflow, which can be applied to multiple biopsy sources and biomarkers, yielding data of comparable quality.

From this workflow, a large amount of standardized data can be obtained. For each patient, between 34 and 194 cells, depending on the burden of the disease, were imaged semi-automatically; per cell, a minimum of 350 tracks and a total of more than 7 million tracks were obtained.

### Machine learning-based data analysis

The large single-molecule datasets generated required a specialised pipeline for analysis to: i) extract the maximum number of features to describe the nanoscale organization and behaviour of the biomarkers; ii) visualize this information-rich dataset at the molecular-, cellular- and patient-level to be able to formulate testable hypothesis about the underlying mechanisms and iii) enable cell classification (e.g. cancer versus healthy cells).

Our group and others developed robust tools to extract mobility parameters from PAINT-SPT data^12,13^, showing how diffusion reflects shifts in functional states^15,38,39^ and that mobility parameters constitute a fingerprint that can be used to classify cell types. Here, we used the deepSPT platform, a machine-learning platform recently developed by Hatzakis and coworkers, to extract a wide range of mobility features and classify diffusional states, an overview of the parameters is given in Table S3.^17^

First, we segmented the cells in the field of view (Fig. 3Ei), followed by deepSPT analysis, for which a custom hidden Markov Model (HMM) was trained. Such a model predicts different states of mobility present within each track. To have a robust and controlled training dataset, we performed PAINT-SPT with 4 different AML cell lines imaged with the same probe that was used for the patients (Fig. 3Eii). In total 200 cells with a combined number of more than 600.000 tracks were used to train the model. The most consistent, accurate and interpretable configuration resulted in a three-state model, corresponding to free diffusion, oligo/dimerization, and receptor immobilization (Fig. S5).

Overall, our analysis pipeline generates over 40 features, including: i) the density of molecules per area of the cell; ii) the most probable diffusion mode (e.g. Brownian, confined, anomalous, directed); iii) a set of diffusion-related parameters describing speed, confinement, heterogeneity and iv) diffusion states and their changes within a track. These diffusional and temporal fingerprints together result in a multiparametric dataset that provides a detailed description of receptor behaviour across molecules, cells and patients (Fig. 3F-H).

### Quantifying receptor abundance and heterogeneity

We started comparing a cohort of AML patients for their molecular-level organization of biomarkers. The biopsies were taken for diagnostic purposes (A samples), and for four patients, an additional sample was taken later in the treatment (B samples). First, we analyse the variations in track density, which serves as a proxy for receptor abundance on the cell membrane. Note that single-molecule detection ensures a better sensitivity in estimating receptor density in samples with very low but non-zero expression^40^ and provides a direct and quantitative estimation (molecules per membrane area) of expression, going beyond the positive-negative dichotomy.

Even at a visual inspection, differences between patients are apparent (Fig. 4A). We observed patients with high (e.g. 3A, left), intermediate (e.g. 2A middle) and low (5A, right) expression present in the dataset. Note that we see patients with 0 tracks per cell, but we can detect ultra-low expressions down to 0.01 tracks/μm^2,^, which roughly translates to a few tens of receptors per cell.

**Figure 4:**
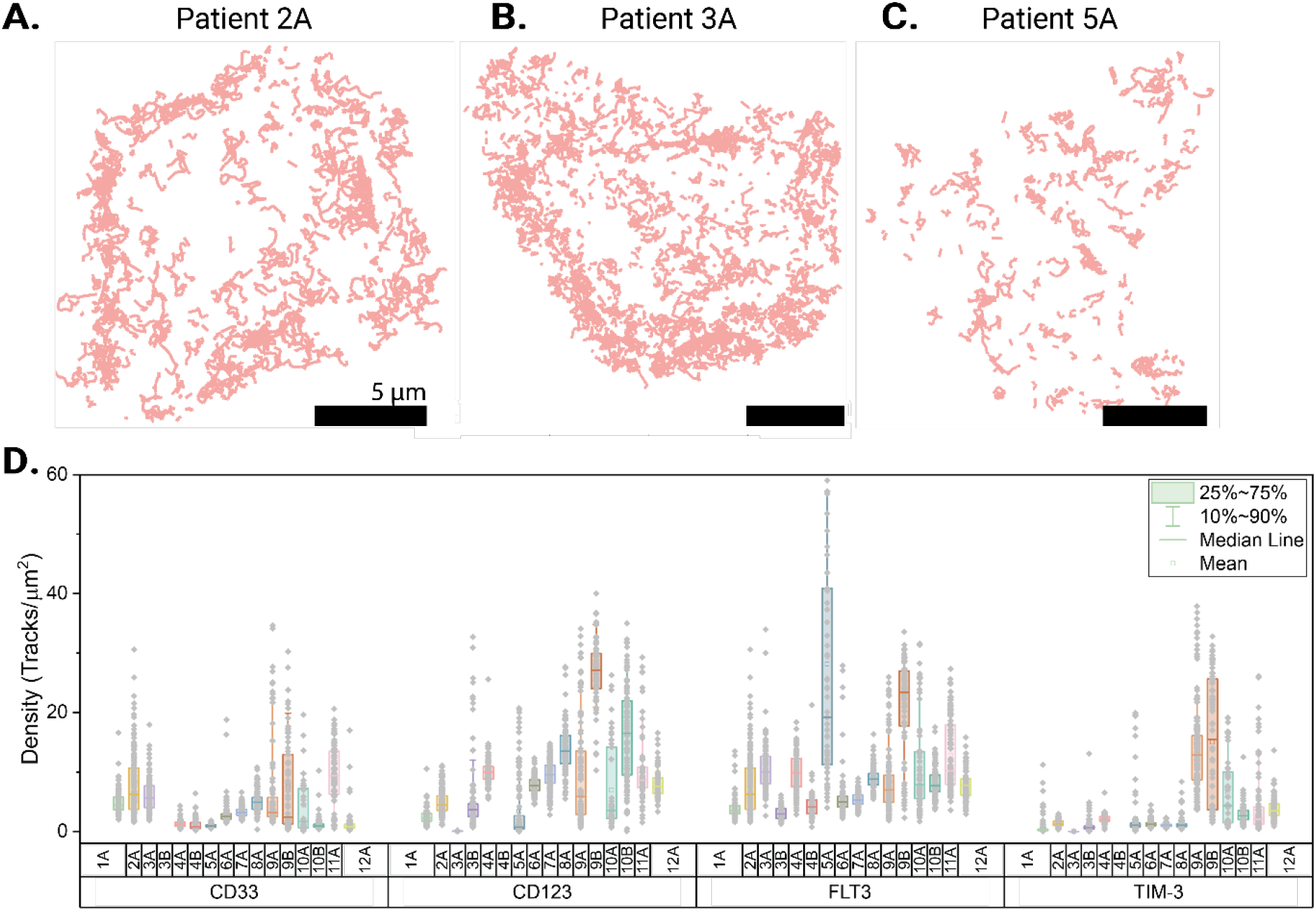
Density of receptors in clinical samples. A-C) Representative images of tracking of CD33 on AML cells of patient 2A, 3A and 5A, respectively. D) The receptor density of each patient is based on the density of tracks/μm^2^ per cell. Some columns are empty because there was not enough sample to measure. Minimum number of cells per patient, per receptor: 60, except for patient 10A (minimum 35).

Fig. 4B shows the receptor density for all the biopsies measured for the four biomarkers of interest: FLT3, CD33, CD123 and TIM-3. As expected, receptor expression varies significantly between patients. Beyond patient-to-patient variability, we assessed intra-patient heterogeneity, using the coefficient of variation (CV) calculated for each receptor (Fig. S6). Some patients or receptors exhibit high CVs, suggesting notable intra-tumour heterogeneity, while others show more uniform expression.^41^

Interesting observations can be made about changes in receptor expression post-treatment and their correlation with the clinical data of the patients (see SI for an extensive discussion). In particular, FLT3 and CD123 vary after treatment, while TIM3 and CD33 are more stable. Notably, high CD123 in NPM1-mutant patients, such as patients 3 and 4, has been linked to improved outcomes when treated with venetoclax plus hypomethylating agents, providing potential therapeutic guidance.^42^ Receptor expression is currently not monitored over the course of treatment, which would be made possible with our approach and could potentially give insights in treatment resistance or changes in subpopulations. Usually, FLT3 expression is upregulated in AML and not in the myelodysplastic syndrome (MDS), which is supported by the low expression levels found for patient 6, who suffers from MDS.

### Receptor mobility reveals patient-specific signatures

Next, mobility fingerprints were compared within the AML patient cohort. To the best of our knowledge, membrane biomarker single-molecule dynamics has never been measured in patients, raising three important questions: i) do patients present a unique mobility fingerprint, or is there a common nanoscale organization and behaviour? ii) how do biomarker dynamics relate to other clinical data, such as genomic profile? iii) are these fingerprints different and predictive in cancer and healthy cells?

To define and visualize the dynamic fingerprint of individual patients, we first combine all the mobility features obtained from deepSPT into a Uniform Manifold Approximation and Projections (UMAP) visualization. Such analysis, common in genomics and proteomics, allows plotting high-dimensional data. Enabling of visualization of differences in mobility patterns between different patients, cells and individual molecules (Fig. S7). For track-level UMAPs, a high correlation between clustering of tracks and their diffusive behaviour was found (Fig. S8). UMAPs per cell are shown in Fig. 5A for FLT3 and for the other markers in Fig. S9. These UMAPs show individual cells as dots that are positioned in the UMAP space depending on the mobility features (averaged per cell).

**Figure 5:**
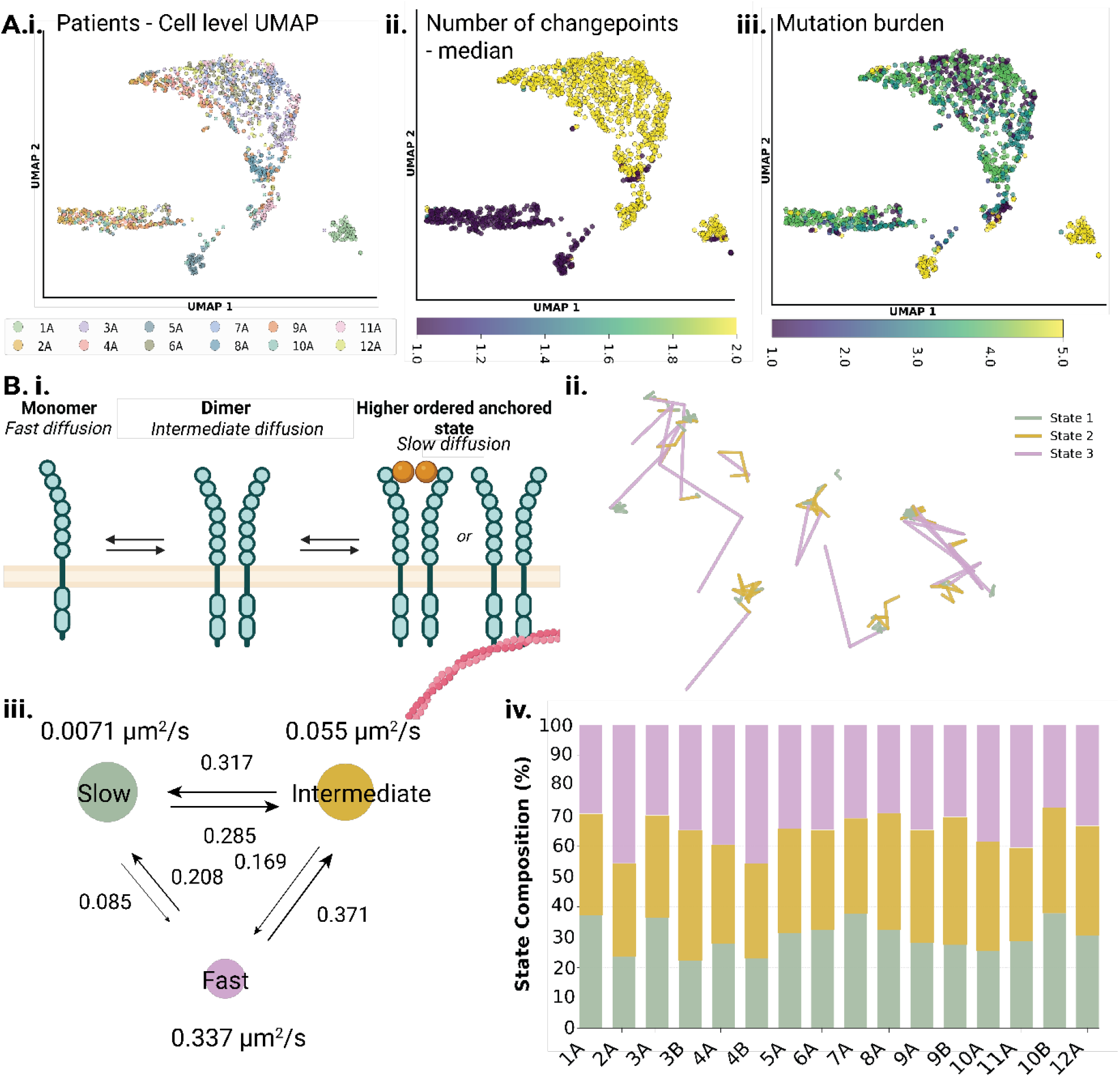
In-depth analysis across patients using UMAP and HMM. A) Cell-level UMAPs of FLT-3 analysed on all AML first biopsy samples (A) colour coded by i) patient number, ii) the median number of change points in a track iii) mutation burden. B) HMM results for all patients. i) schematic of the theoretical behaviour of FLT-3 in which a monomer freely diffuses, a dimer has intermediate mobility and an activated dimer has slow diffusion. ii) Examples of tracks that contain switches. iii) HMM model results with the predicted speeds per state and the switching probability. iv) Average state composition per patient.

Fig. 5Ai shows the UMAP coloured by patients, showing few key insights. First, some patients (1A and 5A) show a very compact distribution, far from the main cluster, indicating different behaviour of these patients. The other patients are closer to each other, but remain in compact clouds per patient, meaning that cells from the same patient are more similar to each other than to those of another patient (Fig. S10). Sometimes, one patient shows two populations of cells, indicating intra-patient heterogeneity. A similar conclusion can be drawn for the other receptors (Fig. S9). All in all, these results show that patients have a unique fingerprint of receptor mobility that can be unveiled for the first time with PAINT-SPT analysis and for specific patients, the difference is very substantial.

After establishing that patients display a unique mobility fingerprint, we investigate the biological reasons for these differences. We found that parameters related to receptor interactions are the drivers of these differences. First, we observed that the number of times a receptor changes behaviour (e.g. from fast to slow due to dimerization) within a single track is one of the key factors that drive the cluster separation in the UMAP (Fig. 5Aii). This is confirmed by the HMM analysis that divides single-molecule tracks into three distinct states that represent different types of diffusive behavior (Fig. 5Bii). For each patient, we can describe both the relative abundance of these states and the likelihood of transitions between them (Fig. 5Biii and iv). Notably, patients 2A and 4B show a higher proportion of freely diffusing, monomeric FLT3 receptors compared to the other patients in this cohort. Also, the confinement ratio changed significantly between patients (Fig. S11). While this is not studied for FLT3, there is an established biological framework for RTKs, where low confinement subdiffusive behaviour relates to homo-interactions of the receptor (e.g. oligomerization or multimerization) and interactions with signalling platforms (e.g. ordered lipid domains, heterodimerization with co-activators, recruitment of intracellular effectors). So overall, we conclude that patients’ differences are mostly due to the tendency of biomarkers to activate and interact with signalling partners.

We associate the mobility fingerprints with standard clinical diagnostic practice of AML patients such as mutation burden. We hypothesized that mutations could result in a different network of interactions and localization, and this would be reflected in receptor mobility. Figure 5Aiii correlates the mobility UMAP with patient mutations. It shows the separation of patients with a high mutation burden (5 mutations) from patients with a lower mutation burden. While the small number of patients of this study prevents any extensive conclusion, the observed trend is interesting for future work.

In this context, our measurements can detect nanoscale organization and interactions showing that biomarkers have different molecular behaviour in different patients, capturing not only expression levels, but also behavioural phenotypes that may reflect underlying functional heterogeneity.

Finally, we moved to use the mobility fingerprint for patients’ classification. First, we cluster patients in sub-classes with a dendrogram analysis. This approach clusters the samples based on both similarity between patients and between features. In Fig. 6A and S12, the dendrogram of the FLT3 receptor is shown, which reveals interesting relationships between patients. Three main groups can be observed: i) patient 2A, 5A and 10A cluster together and are different from all the other patients, the other patients further divide in ii) 4A, 6A, 9A and 12A and iii) 1A, 3A, 8A, 7A and 11A, with again internal division between patients. This seems to indicate that patients have a unique mobility fingerprint, but categories based on similarity start to emerge. An interesting future perspective of this is the expansion of this study to a larger cohort of patients to make robust conclusions on mobility sub-types in AML disease and their relations with therapy outcome.

**Figure 6:**
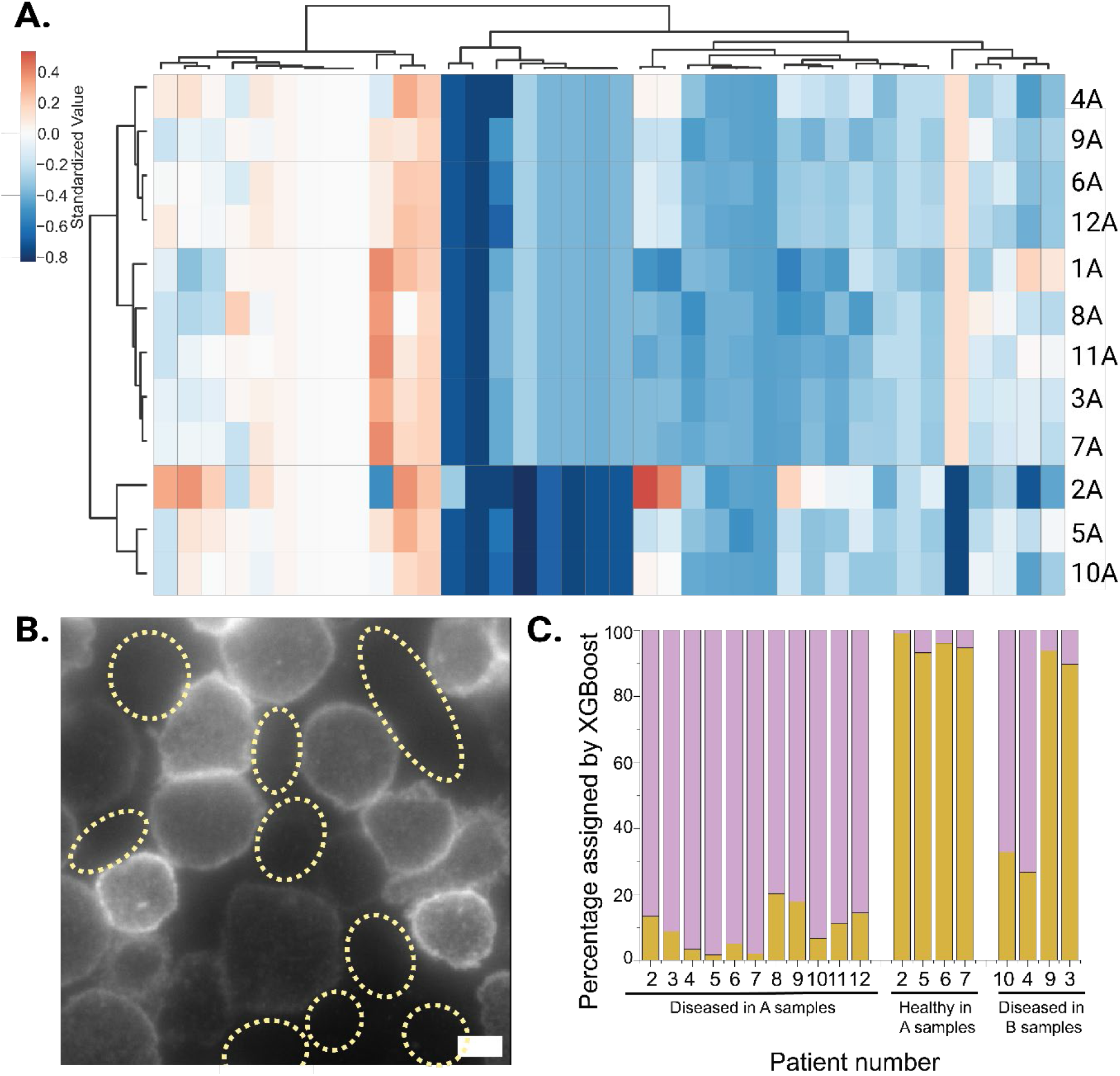
Patient stratification and classification. A) Dendrogram analysis of all DeepSPT results. B) Example of a field of view with a mix of diseased and healthy cells, healthy cells indicated with yellow dotted circles. C) Percentage of cells assigned to healthy and unhealthy per sample by the XGBoost model.

Next, we analysed if the single-molecule fingerprint allows to distinguish healthy from cancer cells. Within a single field of view of all patients, both CD34/CD117-positive and negative cells can be distinguished, as indicated in Fig. 6B, serving as ground truth for healthy and cancer cells. We employed an eXtreme Gradient Boosting (XGBoost) supervised learning strategy to classify our patients (ROC curves can be found in Fig. S13). As is demonstrated in Fig. 6C, the model for FLT3 correctly predicts >80% of the cells to be healthy or diseased. Interestingly, this accuracy lowers for the B samples, where the patient is already being treated, indicating changes of the cells towards the healthy phenotype. Moreover, not all biomarkers are predictive, with FLT3 being the most suitable to identify the cancer cells. To the best of our knowledge is the first time cancer is identified by molecular mobility and behaviour rather than by their expression underscoring the potential of PAINT-SPT for classification of the disease state of patients.

### Comparison of models for nanoscale membrane organization

The primary biopsy site (bone marrow) is the real setting to assess patients’ cells with diagnostic potential. However, this is not always accessible and other cell sources may play a role in preclinical and clinical research such as peripheral blood cells or cell lines. This open the question how representative of the primary source these models are in terms of nanoscale organization and receptor mobility.

To investigate whether *in vitro* models represent receptors on AML cells of patients properly, cell lines were compared with clinical samples. The mobility parameters of both patients and cell lines were plotted in the same UMAP, which is shown in Fig. 7A. The patient cells cluster and are very far from the cell lines, meaning that between patients there are more similarities than between patients and cell lines. An exception is TIM-3 (7Aiii), where the cell lines and patients are all close together, indicating a higher similarity. Together, this shows that although cell lines offer advantages in terms of accessibility and consistency, they may fail to capture the biological complexity of human biology due to a diverse population of mutations and genetic drift^43^. While cell lines are a necessary tool in cell biology, our results strengthen the argument for patient-based analyses to study fundamental mechanisms of disease.

**Figure 7:**
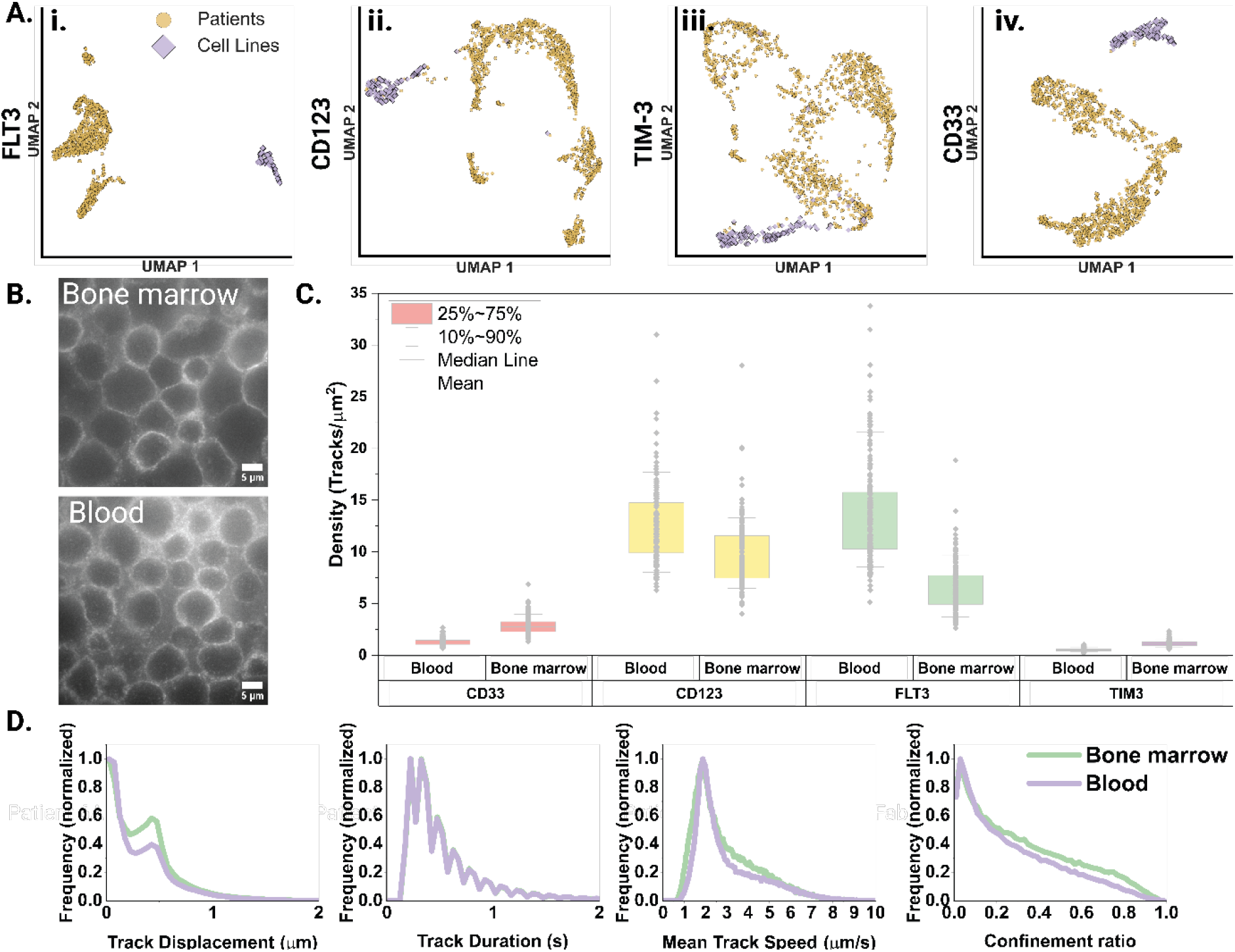
Comparing sample sources. A) Patient cells compared to cell lines. i) FLT3, ii) CD123, iii) TIM-3 and iv) CD33. Orange circles indicates patient samples, purple diamonds indicate cell line cells. C) Patient cells derived from bone marrow compared to peripheral blood. i) Identification step shows high number of cells identified as CD34/CD117 positive in both blood and bone marrow, scale bar 5 μm, ii) expression levels of all receptors between PB and BM. All differences p<0.0001 according to two sample t-test, iii) selection of mobility parameters on FLT-3.

To explore if blasts are similar between different body fluids, we analysed the PB and BM from the same patient. Fig. 7Bi demonstrates that PB-derived blasts were successfully immobilised and identified enabling direct comparisons between bone marrow-resident and circulating blast populations, which potentially differ in subtypes of cells,^43,44^ but show high concordance between mutations.^45,46^

In Fig. 7B-D, receptor surface densities in blood and bone marrow are compared. Densities show small but statistically significant differences, suggesting minor changes in subpopulations between compartments. Moreover, when comparing mobility parameters of these receptors (Fig. 7D and Fig. S14), trends remained very consistent between compartments. These findings show that overall, blood samples are representative of nanoscale organization and provide an accessible way for therapy monitoring.

## DISCUSSION

In this work, we aimed to develop a workflow for live-cell single-molecule imaging on clinical liquid biopsies able to overcome the challenges and restrictions attached to use of clinical samples.

We showed that our workflow meets the requirements of clinical samples, ensuring cell of interest enrichment, identification and viability. Using Fab domains as probes, we performed PAINT-SPT without compromising resolution, sensitivity and data quality and we achieved sufficient throughput through microscope automation.

To validate our workflow and show the generality of this approach, we imaged multiple biopsies spanning two types of cancer (AML and lung cancer), three biopsy types (bone marrow, blood and pleural effusions) and multiple biomarkers. For each patient, tens to hundreds of cells were measured and a one-of-a-kind dataset composed of over 7 million detected single-molecule trajectories has been obtained and analysed with a variety of AI and statistical tools.

By focusing on mobility parameters (*e*.*g*. displacement, diffusion coefficient, confinement) and diffusive behavioural changes, we move beyond traditional phenotyping based on surface marker intensity. This approach provides a new dimension to biomarker analysis that has proved very effective for understanding patients’ biomarker properties at the nanoscale as well as for patients’ classification.

To advance this method towards clinical applications, we envision three perspectives: i) Larger patient groups. More patients are needed to solidify conclusions and ensure statistical robustness. The genetic profiling framework in AML provides a valuable starting point, but larger patient groups with overlapping genetic profiles and from different diseases are necessary to determine the true diagnostic potential of receptor mobility and density measurements. Additionally, the treatment outcomes can give valuable insights into the predictive potential of our approach. ii) Mechanistic understanding of receptor mobility. To move beyond descriptive readouts and speculations, mobility patterns should be linked to a biological function. This can be done with complementary assays in biopsies and cell lines. iii) Integration with clinical and genomics data. Single-molecule measurements can be further crossed with established diagnostics such as genetic mutations, karyotyping and immunohistochemistry, as well as correlated with clinical outcomes. In this work, initial steps were taken, but they should be deepened as the number of patients increases. Establishing these connections will be key to demonstrating the diagnostic power of nanoscale receptor profiling.

In conclusion, our work provides a robust method for clinical studies of biomarkers at the single-molecule level and shows its potential to identify unique patients’ nanoscale phenotypes in health and disease. We envision that PAINT-SPT represents a valuable tool for biomarker analysis and support in future clinical applications such as diagnostics, personalised cancer therapy, functional drug-screening ex vivo and therapy monitoring based on liquid biopsies.

## ONLINE METHODS

### Ethics statement

This study was conducted in accordance with the ethical principles outlined in the Declaration of Helsinki. Ethical approval for the non-WMO study for AML was obtained from the Board of Directors of Catharina Hospital Eindhoven (Approval Number: nWMO-2024.078) and all procedures involving human participants were reviewed and approved before the commencement of the study. All participants provided written informed consent before their inclusion in the study. They were fully informed about the purpose, procedures, potential risks and benefits of the research. Participation was voluntary and participants were given the option to withdraw at any time without any consequences. Confidentiality and anonymity of the participants was ensured through secure data handling and storage, with all personal identifiers removed or anonymised in the final dataset. The researchers adhered to all relevant institutional and national guidelines for research involving human participants in a nWMO study. Regarding the pleural effusion samples, all samples received were anonymized and no information about patient details was known to the researchers performing the microscopy. All National and European legislation concerning data privacy was followed.

### Digestion of antibodies

The antibodies used for SPT were digested using a Pierce Fab Micro Preparation Kit (ThermoFisher). Briefly, the papain solution was washed with Fab Digestion buffer, after which the antibodies in digestion buffer were added to the papain bed. This reaction was incubated for 6 hours, in the ThermoMixer at 37 °C and 700 rpm. The product was obtained from the papain using centrifugation at 5000 g for 1 minute, followed by a wash with 130 μL PBS. The Fab fragments were separated from the Fc domains and the undigested antibodies using a Protein A column. For this, the solution was added to the column and incubated for 10 min at RT and 700 rpm. After centrifugation at 1000 g for 1 min, the flow through was saved and the column was washed with an additional 200 μL of PBS. Fab concentration was measured by absorbance at 280 nm using a Nano-Drop One (Thermofisher). SDS-PAGE was performed to confirm a successful product (Fig. S15).

### Fluorescent labelling of Fab fragments and full antibody

For labelling the Fab fragments, ATTO 643 (ATTO-TEC) was used. For the full Vadastuximab antibody, ATTO 532 (ATTO-TEC) was used. The dye was dissolved in DMSO to reach a concentration of 10 mg/mL. A 80x molar excess of the dye was added to the Fab fragments and the reaction was incubated for 2 hours and 30 minutes under moderate stirring (400 rpm) at 25 °C in a ThermoMixer (Eppendorf). Afterwards, the reaction mix was transferred to a dialysis tube with a 20kDa molecular weight cut-off. Dialysis against PBS was performed overnight to get rid of excess dye and PBS was renewed after the first 2 hours. Protein concentration and degree of labelling were determined using a Nano-Drop One (Thermofisher) with PBS as the blank measurement. Degrees of labelling can be found in Table 1.

**Table 5.1:**
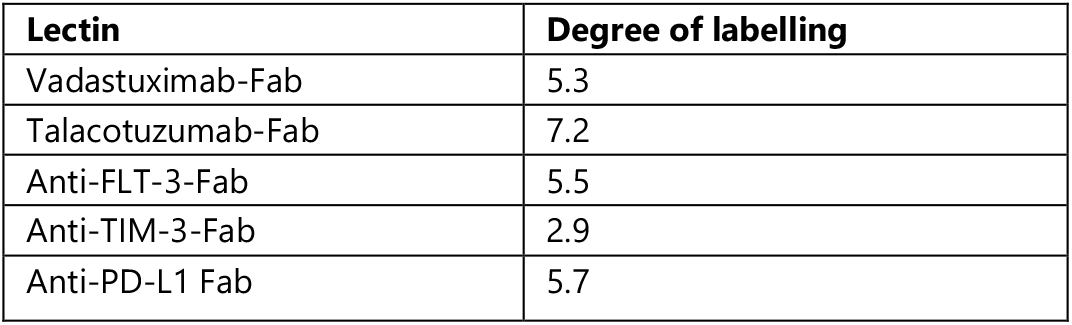
Overview of the probes used in this research and their respective degree of labelling. All Fabs were labelled with ATTO-643, Vadastuximab-Full was labelled with ATTO-532.

| Lectin | Degree of labelling |
| --- | --- |
| Vadastuximab-Fab | 5.3 |
| Talacotuzumab-Fab | 7.2 |
| Anti-FLT-3-Fab | 5.5 |
| Anti-TIM-3-Fab | 2.9 |
| Anti-PD-L1 Fab | 5.7 |

### Retrieving cells from pleural effusion biopsies

Pleural effusion samples from lung cancer patients are provided by the Catharina Hospital in Eindhoven. Circulating tumour cells are isolated through negative enrichment using pluriSpin Human PBMC Depletion (PluriSelect) with the following protocol: pleural effusion samples are filled up to 8 mL with PBS-BSA 0.1%, 40uL of the pluriSpin beads are added and the mixture is incubated at room temperature for half an hour in a tube roller. Then, 2mL of PBMC Spin Medium (PluriSelect) are slowly added to the bottom of the tube without disrupting the interface between the two media. Samples are centrifuged at room temperature and 800g for 15 minutes with acceleration and brake settings off. Density gradient centrifugation separates red blood cells and PBMCs (bound to depletion beads) at the bottom of the tube, while less dense CTCs are collected at the interface (i.e. buffy coat). Then, few millilitres are collected from the interface to ensure all CTCs are recovered and diluted with PBS-BSA 0.1% to 15mL. Tubes are then centrifuged at 300g for 10 minutes, the supernatant is discarded and the pellet is resuspended in CTC medium containing phenol red free RPMI1640 medium (Gibco), 20 ng/mL epidermal growth factor (EGF), 20 ng/mL basic fibroblast growth factor (FGF2) and B27 supplement.^47^

For positive enrichment, the pluriBead Human CD326 cell separation kit (pluriSelect) was used to isolate EpCAM positive cells using the provided protocol.

### Retrieving cells from bone marrow aspirate or peripheral blood of AML patients

From the AML samples received from the hospital (between November 2024 and August 2025), the red blood cells were removed by adding 4 mL ammonium chloride to 1 mL of bone marrow aspirate or, in the case of peripheral blood, 9 mL of ammonium chloride to 1 mL of blood. This solution was then left on ice for 10 min, after which the white blood cells were obtained by centrifuging 3x at 150g for 5 min, replacing the supernatant with RPMI media supplemented with 10% FBS and 1% penicillin/streptomycin (100 U/mL and 100 μg/mL respectively). After the last step, the cells were resuspended in 400 μL of RPMI.

### Optical setup

PAINT images were obtained using an Oxford Nanoimager Microscope (ONI), prewarmed to 37 °C. TIR conditions were used for captured cell PAINT experiments, temperature and laser powers of 488nm and 640nm lasers were kept constant (20 and 40 mW respectively). Fluorescence was recorded using an x100/1.4-numerical aperture oil immersion objective, passed through a beam splitter. Emission was detected in multiple 428 x 500 μm ROIs.

### Sample preparation captured LC cells

To capture lung cancer CTCs, slides were coated with RGD to promote cell adhesion. For this, a µ-Slide VI 0.5 Glass bottom slide (Ibidi) was treated with UV/Ozone for 10 minutes, after which PLL-PEG RGD 3-10% (SuSoS) was added at a concentration of 0.1 mg/mL. This was left to react for 15 min. Nonreacted PLL-PEG was removed by washing three times with PBS. Then, 40 μL of the purified clinical sample was added to the prepared chamber. The cells were left to incubate for 1 hour in the incubator, after which non-bound cells were washed off with prewarmed CTC medium.

### Sample preparation captured AML cells

To capture AML cells, slides were prepared with capture antibodies. For this, a µ-Slide I 0.6 Luer Glass bottom slide (Ibidi) was treated with UV/Ozone for 10 minutes, after which PLL-PEG biotin 50%/50% (SuSoS) was added at a concentration of 0.2 mg/mL. This was left to react for 15 min. Nonreacted PLL-PEG was removed by washing three times with PBS. Neutravidin (Fisher Scientific) was added to the slides at a concentration of 0.2 mg/mL and again incubated for 15 min, followed by three washes with PBS. A mixture of biotinylated-CD44 (IM7, Fisher Scientific) and biotinylated-CD45 (BAM1430, Biotechne) antibodies at a concentration of 0.1 mg/mL was prepared and added to the slide and left for 15 min, followed by two washes with PBS and one wash with RPMI media. 100 μL of the purified clinical sample was added to the prepared chamber. The cells were left to incubate for 45 minutes in the incubator, after which non-bound cells were washed off with prewarmed Leibovitz imaging media (Invitrogen) containing 10% FBS.

### Single-molecule imaging of the LC receptors with PAINT

Before imaging, cells were incubated with 1:20 dilution of anti-hEpCAM Alexa Fluor 488-conjugated antibody (Biotechne) in CTC medium for 10 minutes. Afterwards, cells were washed with fresh medium, and medium containing 5 nM of ATTO643 labelled PD-L1 Fab was added to fill the channel and reservoirs of the slide. Imaging was started immediately afterwards on the Oxford Nanoimager microscope (ONI), prewarmed to 37 °C. Using brightfield illumination and a 488nm laser, EpCAM-positive cells were located, and PD-L1 Fab binding was imaged for 2,000 frames at a rate of 33.3 Hz with a 640nm laser. In addition, a brightfield image and one frame of the reference EpCAM antibody channel were also acquired prior to acquiring the Fab fragment images.

### Single-molecule imaging of the AML receptors with PAINT

Before imaging, the cells were washed thrice using a prewarmed Leibovitz media supplemented with 10% FBS. Afterwards, an imaging solution mix containing 4 nM probe plus 100x diluted identification antibodies (CD34 and CD117) was added to the wells. For the cell lines, these latter antibodies were not used. Imaging was started immediately afterwards on the Oxford Nanoimager microscope (ONI), prewarmed to 37 °C. Data was collected for 6,000 frames at a rate of 33.3 Hz, with a total acquisition time of 5 min and a minimum number of cells per probe of 100. For cell identification, 5 frames were taken of the identification antibodies prior to acquiring the Fab fragments.

### Analysis of PAINT

Acquired movies were analysed with the NimOS software (1.18) to reconstruct the binding events of Fab probes by fitting a two-dimensional Gaussian to individual fluorescence spots with a localisation precision of at least 30 nm. Within the software, single particle tracking (SPT) can be performed to obtain the trajectories of Fab probes bound to receptors in consecutive frames. The following parameters were used for AML: maximum frame gap, 3; maximum distance between frames, 0.5 μm; exclusion radius, 1.0 μm. A filter for a minimum number of steps of 5 and a minimum diffusion coefficient of 0.05 μm^2^/s was applied. For LC, the following parameters were used: maximum frame gap, 3; maximum distance between frames, 0.3 μm; exclusion radius, 0.8 μm. A filter for a minimum number of steps of 5 and a minimum diffusion coefficient of 0.04 μm^2^/s was applied. Then, the csvs of the tracking analysis were imported into Matlab R2023b, where ROIs were drawn around the cells that were identified in the identification step, to cut them from the csv.

After an initial analysis of the step lengths of the tracks, an anomaly was found for the cell line data where the density of steps was increasing for steps larger than 0.4 μm. These large steps were caused by tracks in close proximity in time and space to one another, which were erroneously connected by to these large steps. The tracks were separated with a custom python file, which removes steps above 0.4 μm and constructs new tracks by reconnecting the localisations with separated connections. This reconnection was limited to the localisations within a frame gap of 3 frames and a maximum distance of 0.4 μm. To ensure comparability between the cell line and leukaemia datasets, the step length filtering was applied to both.

The density of binding events was calculated by counting the number of trajectories within the region of interest determined by the contour of the cells. Data from at least 100 cells were used for quantitative analysis. To analyse the diffusional behaviour of the tracks, the deepSPT^17^ pipeline was used. The features produced by deepSPT are described in Table S3. In addition to the original features, two new features were introduced: the confinement ratio, which indicates the straightness of a track, and the median photon count of the track. These track fingerprints were then used for downstream analysis. Some features of the original deepSPT were excluded in the follow-up analysis as their value was 0 for all tracks and all patients.

### HMM training, validation and analysis

A custom HMM model was trained to be used within the deepSPT pipeline. The cell line dataset was used to train a general model for receptor diffusion behaviour. Both a three and four state variant of the model were trained using the pomegranate library using the DenseHMM functionality, which were validated in two ways: the output of the cell line models were compared with the output of a model trained on the leukaemia dataset itself and with the Akaike Information Criterion (AIC), which incorporates both the amount of model parameters and the loglikelihood of the model. The resulting accuracies and AIC scores are displayed in Fig. S5. The distribution of the states per patient is found by predicting all steps in the dataset and dividing the number of steps predicted as a specific state by the total number of states for each patient. The random state used to train all the HMMs was 42, which will be used for every other analysis described (when applicable).

### Dimensionality reduction

A Uniform Manifold Approximation and Projection (UMAP) was performed in Python, using the UMAP library. The dataset was first standardised (all the features are transformed to have a mean of zero and a standard deviation of one) using the StandardScaler module from scikit-learn. To create cell-level representations of the diffusion behaviour of the tracks, the mean, median and standard deviation of the track fingerprints were calculated for each cell. Then the UMAP was performed with the hyperparameters number of neighbours and minimum distance set to respectively 15 and 0.1 for the track-level and 50 and 0.05 for the cell-level. Colouring was based on the HDBSCAN cluster for the track-level UMAP and the patients for the cell-level. Both track and cell-level UMAPs were also coloured by specific metrics, as indicated in Figures 4 and 5.

### Clustering

The HDBSCAN clustering was performed using a min_cluster_size of 5% of the total data and min_samples set to 1 on the scaled data. The adaptive cluster size was used to prevent small noisy clusters, while still allowing the algorithm to find the optimal clusters. The lower min_samples allows for more datapoints to be assigned a cluster instead of being treated as noise. For the other hyperparameters, the default settings were used. As the high number of tracks severely impacted the computational time of the algorithm, the GPU-accelerated HDBSCAN algorithm from the cuML library was used. However, the functionality and output should be equal to the HDBSCAN library if no GPU is available. Hierarchical clustering analysis was performed to identify patient groupings based on the median of the scaled DeepSPT fingerprint data. Hierarchical clustering was performed using Ward’s linkage method with Euclidean distance metrics on the standardized feature data. This was done with the linkage, pdist and fcluster functions from scipy. Clustered heatmaps were generated using seaborn’s clustermap function. The dataset was limited to the untreated (A) patient samples for this analysis to find both inter- and intrapatient level differences and similarities in track behaviour.

### Cell classification

Classification on the cell level was also performed. With the same mean, median and standard deviation aggregation as described previously, the cells of patients with a clear distinction between healthy and diseased cells in the anti-CD34/CD117 staining were used to train XGBoost classifiers. Only four patients met this requirement, which were each used once as a test sample, while the others were being used as training data (leave one out cross-validation). The XGBoost used the following parameters: max_depth, 3; learning_rate, 0.1; n_estimators, 100; subsample, 0.8; colsample_bytree, 0.8; and the scale_pos_weight was calculated to correct for the class imbalance between healthy and unhealthy. The data scaling was done independently for each fold by fitting the scaler only on the training data and then applying it to the test set to avoid data leakage between the train and test sets. The models trained for each fold were then used to predict the amount of healthy and unhealthy cells in each sample in the complete dataset.

## DATA AVAILABILITY

Data is accessible upon reasonable request (due to privacy rules).

## CODE AVAILABILITY

All codes for SPT are available at Github N4N.

## ACKNOWLEDGEMENTS

Schematic illustrations in all figures were created using Biorender.com. The authors would like to thank Anna Świetlikowska for performing the SDS-PAGE of the Fab domains and the Department of Special Haematology of the Catharina Hospital for collecting the samples used in this study. M.M.E.T., R.R. and L.A. thank the financial support from The Netherlands Organization for Scientific Research (NWO VIDI Grant 192.028 and OCENW.XS24.2.147). R.R. and L.A. thank the European Research Council for financial support (no. ERC-2022-POC2 - 101100873 - NANODIAGNOSTIC).

## AUTHOR CONTRIBUTIONS

Conceptualization was done by R.R., V.S., M.J.C., D.v.d.K., M.M.E.T. and L.A. Patient recruitment was done by M.J.C., V.S., D.v.d.K., H.C. Wet lab methodology was formulated by R.R., M.M.E.T., F.C., M.E.M. and L.A.. All measurements on patient cells were performed by R.R. and M.M.E.T. Data analysis strategy was formulated by R.R., M.M.E.T., T.H.W.H., M.V. and L.A. Data analysis was performed by R.R., M.M.E.T. and T.H.W.H. The original draft was written by M.M.E.T. and L.A. in consultation with all other authors.

## COMPETING INTERESTS

The authors declare no competing financial interest.

## Notes

### Competing Interest Statement

The authors have declared no competing interest.

