## Supplementary information for "Single-molecule imaging and tracking on clinical liquid biopsies reveals cancer biomarkers nanoscale organization and heterogeneity"

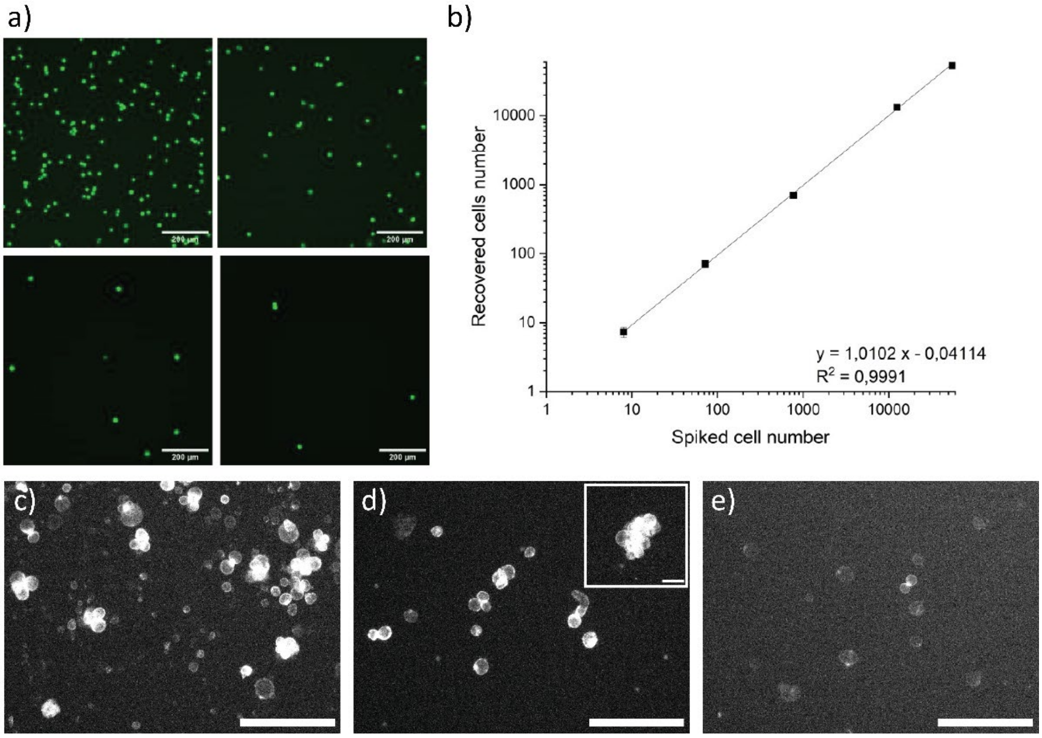


**Figure S1: Comparison of positive and negative enrichment.** A) fluorescently labeled MDA-MB-468 cells in blood from a patient. Scale bar: 200 µm B) Graph of the recovery rate of spiked cells into patient material. C) Microscopy image of fluorescently labeled cells within a pleural effusion sample (anti-EpCAM) before processing. D) EpCAM labelling of CTCs after negative enrichment and E) after positive enrichment. Scale bar: 100 µm.

By spiking fluorescently labelled cancer cells to immune cells to simulate a pleural sample, a recovery rate of 99% was found with the proposed negative enrichment protocol. With an EpCAM-based positive enrichment, a reduction in EpCAM in CTC was found, suggesting that removal of the cancer cells from the beads could result in damage to the cells or loss of the identifying biomarker. Therefore, we chose to optimize the negative enrichment using peripheral blood mononuclear cells (PBMCs) depletion beads followed by density centrifugation, which already removes RBCs and peripheral blood polymorphonuclear cells (PMNs), effectively removing all red and white blood cells and leaving the lung cancer circulating tumour cells untouched. Fig. 2B shows how negative enrichment removes most of the immune cells (small size, EpCAM negative) while lung cancer cells are maintained (large size, EpCAM positive). Fig. S2 shows the viability of cancers cells after the enrichment process, and although there is small decline at longer times, they remain viable for many hours.

Viability of MDA-468 and CTCs after purification on EpCAM capturing slides

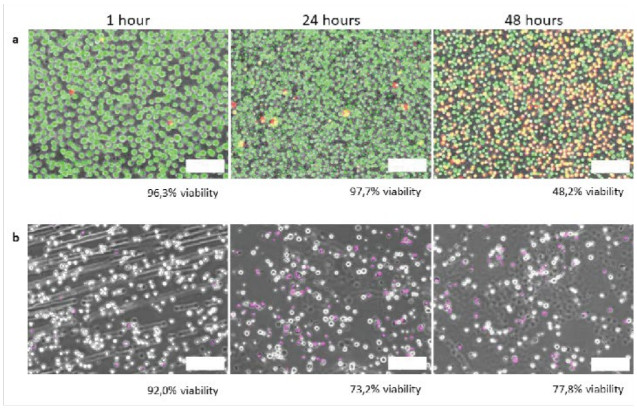


Viability of MDA-468 on EpCAM capturing with and without purification

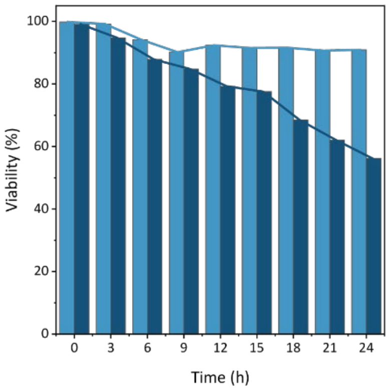


**Figure S2: Viability of MDA-MB-468 and pleural effusion CTCs 24 hours after capture on an anti-EpCAM covered slide.** A) Images of viability comparison at 1h, 24h and 48h for both CTCs and MDA-MB-468 cells. B) Comparison of MDA-MB-468 cells spiked in patient material and purified using negative enrichment and MDA-MB-468 cells captured directly.

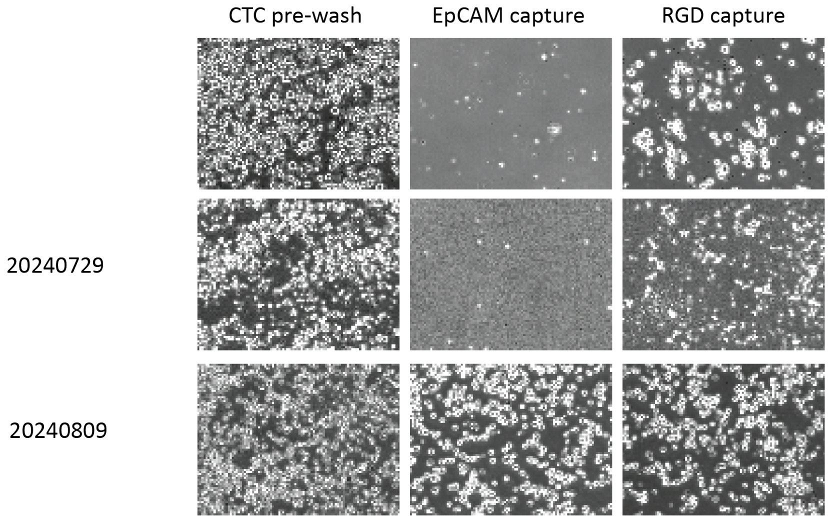


**Figure S3: Comparison of anti-EpCAM capture and RGD capture for multiple patients**.

**Table S1: Karyotyping of the AML patients used in this study**

| Patient | Subtype | Karyotype | Mutations |
|---|---|---|---|
| 1 | AML with myelodysplasia-related changes | XX, 46 | BCOR+<br>DNMT3A+<br>IDH2+<br>SRSF2+<br>STAG2+ |
| 2 | MPAL (AML + ALL), AML with myelodysplasia-related changes | XY, 46 | FLT3-ITD+<br>IDH2+<br>RUNX1+<br>SRSF2+ |
| 3 | AML not otherwise specified | XX, 46 | DNMT3A<br>IDH1+<br>NPM1+ |
| 4 | AML with myelodysplasia-related changes | XY, 46, -7 | NPM1+<br>ASXL1+<br>CBL+<br>GATA 2+ |
| 5 | AML with myelodysplasia-related changes | XX, 47 der (1;19)(q10p;10p) | ASXL1+<br>DNMT3A+<br>TP53 (VAF 1.1%)<br>RUNX1+<br>SETBP1+ |
| 6 | Myelodysplastic syndrome | XY, 46 | Normal |
| 7 | Acute myeloid leukaemia | XY, 46 | FLT3-ITD+<br>IDH2+<br>RUNX1+<br>SRSF2+ |
| 8 | AML with myelodysplasia related gene mutation | XX, 46 | ASXL1+<br>IDH1+<br>SRSF2+<br>STAG2+ |
| 9 | Acute myeloid leukaemia | XY, 46 | ASXL1+<br>GATA2+<br>DNMT3A+ |
| 10 | Starting relapse of AML | XY, 47 +13 | FLT3-TKD+<br>RUNX1+ |
| 11 | Myelodysplastic syndrome/acute myeloid leukaemia | Complex profile with del3p, del5q, del7q, del12p | TP53 (VAF 80%) |
| 12 | Relapsed AML. Previously treated with azacitidine and venetoclax. Previously had a FLT3-ITD, now not detectable. | XY, 46 | CSF3R+<br>IDH2+<br>RUNX1+<br>SRSF2+ |

**Table S2: Immunohistochemistry results of the AML patients included in this study. Last column represents the amount of staining observed in the identification step of the SRM.**

| Patient | Microscopy findings | Immunohistochemistry | Observed CD34/CD117 |
|---|---|---|---|
| 1 | Hypercellular for age (95% cellularity)<br>Presence of blasts with discrete nucleoli<br>Background shows pre-existing erythropoiesis<br>No notable mitotic activity<br>Possible mild nuclear abnormalities<br>Megakaryocytes possibly show detached nuclear lobes<br>Laguesse staining shows no convincing myelofibrosis | MPX: positive staining in blasts<br>CD61: stains megakaryopoiesis with focal microforms<br>CD34: stains clustered blast cells, focal > 20% | Not measured |
| 2 | Hypercellular (90% blast cells)<br>Strong and diffuse expression of TdT and CD34<br>Some cells express PAX5, CD20 and CD10<br>Myeloperoxidase shows scattered positive cells also expressing CD117 | TdT, CD34: strongly and diffusely positive.<br>PAX5, CD20, CD10: positive on some blasts<br>MPX and CD117: scattered positive cells | High degree of staining |
| 3 | Hypercellular (70%) with monotonous appearance and presence of blasts.<br>No myeloid maturation<br>Scattered megakaryocytes with hyperlobulation and detached lobes<br>No notable erythropoiesis<br>No increase in lymphocytes or plasma cells<br>*B sample:* no increase in blasts | No IHC due to correlation with aspirate and FC (CD117 partially positive, CD34 negative) | Weakly present |
| 4 | Hypercellular (89-98%)<br>Oval-shaped cells with moderately defined amphophilic cytoplasm.<br>Morphologically, blast cells<br>Myelopoiesis shows suboptimal maturation<br>Erythropoiesis is minimally present.<br>Megakaryocytes show multiple microforms, including dysplasia<br>*B sample:* not representative | CD43: positive in oval-shaped cells<br>CD68: occasional histiocytic cells stain positive<br>MPX: negative<br>CD34 and CD117: partial staining<br>CD61: positive in megakaryocytes<br>CD34: >5% <10% | Weakly present |
| 5 | Cellularity ~30%<br>Left shift, with immature cells in myeloid lineage that do not mature properly.<br>Erythroid series is arranged in erythrones with some variation in nuclear diameter<br>Myeloid-to-erythroid (M:E) ratio is approx. 3:1.<br>Only a few megakaryocytes are identifiable | CD61: microforms<br>CD34 and CD117: extensive positivity (~30%)<br>CD3: slight T-cell infiltrate<br>CD20: few B-cells<br>CD138: scattered plasma cells with normal distribution | High degree of staining |
| 6 | Hypocellular marrow (10%)<br>Few megakaryocytes present, no notable abnormalities<br>No clustering or microforms<br>M:E ratio within normal limits<br>Myelopoiesis does not mature (MPX negative)<br>Erythropoiesis located in erythron nests, partly contacting trabeculae<br>No morphological synchrony | CD34: shows increase in blasts, estimated around 20%<br>CD117: blasts are mostly negative | High degree of staining |

| | | | |
|---|---|---|---|
| | No increase in lymphocytes or plasma cells<br>No fibrosis | | |
| 7 | Almost completely taken over by uniform blast cells.<br>Only a few erythroids in the background. No additional testing was performed. | | High degree of staining |
| 8 | Hypercellular bone marrow for age (90%) with 70% infiltration by an acute myeloid leukemia.<br>In the background some plasmacytosis is observed with enhanced kappa/lambda ratio (5/1). | MPO+<br>CD117+<br>CD34+ | Few cells with high degree of staining, others weakly stained. |
| 9 | Cellularity is around 80%. The marrow is largely occupied by an immature, blast-like cell population arranged in large fields. In between, there is still some evidence of maturation in the myeloid lineages<br>*B sample:* aspirate 33% blasts, CD34+, biopsy: 20% blasts, CD34+ | MPO+<br>CD33+<br>No recognizable residual erythroid lineage<br>CD61 shows a single megakaryocyte without obvious dysmorphic features.<br>CD34 could not be reliably assessed due to technical problems of unknown cause.<br>The blast percentage is certainly >20%.<br>Same applies to CD117, at least partly positive. | Fluctuates, high in FLT3 and TIM3, lower in CD123 and CD33. |
| 10 | Mildly hypercellular bone marrow. CD61 shows a slight presence of megakayopoiesis with a few microforms.<br>MPX shows an estimated M:E ratio of 1:1<br>MUM1 shows staining of plasma cells, mostly located pericapillary, polyclonal for kappa and lambda with a ratio of 2:1.<br>Suspicion of dysplasia of megakaryopoiesis and possibly erythropoiesis as well.<br>*B sample:* aspirate: 1.6% CD34+ blasts, biopsy: no increase in blasts, few CD34+ blasts found. | CD3+ T lymphocytes<br>CD20+ sporadically a B lymphocyte<br>CD34 shows staining of blast cells, increased in number and partly clustered. The percentage is estimated at >10% and <20%. The blasts do not stain with CD117, although they express MPX. | Low to no staining. Higher with B sample |
| 11 | The intertrabecular bone marrow is hypercellular for age (approx. 90%).<br>The megakaryocytes are increased in number and show dysplastic hyperchromasia; some are small. No clustering observed.<br>Erythropoiesis (CD71) is relatively normally represented compared to myelopoiesis and shows megaloblastic features, with partial contact with bony trabeculae, suggestive of dysplasia.<br>A few well-matured neutrophilic granulocytes (MPX) are seen, although myelopoiesis appears to show impaired maturation.<br>No fibrosis. | CD61: increased in number<br>CD71: normal<br>CD34 and CD117: difficult to interpret pattern (overly strong antigen retrieval). Blast 10%<br>No increase in CD3, CD20 and CD138 | Weakly present |
| 12 | Very hypercellular bone marrow, with a hematopoietic tissue-to-fat tissue ratio of 95/5.<br>Megakaryocytes are largely normal in shape. The increased cellularity is the result of infiltration by myeloblasts. | MPX and CD34: myeloblasts approx. 20%<br>CD117+ | Few cells with high degree of staining, others weakly stained. |

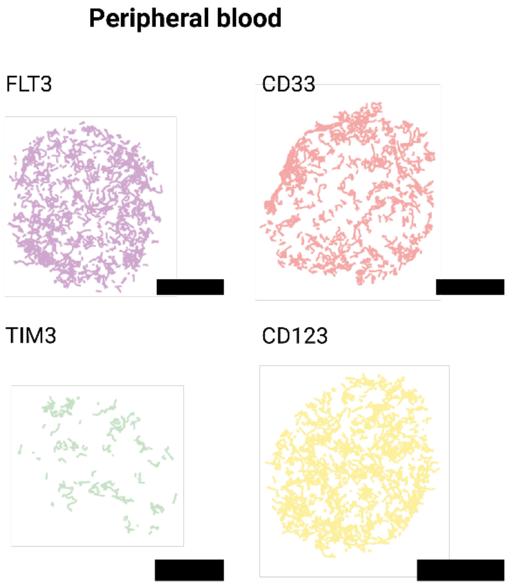


**Figure S4: Probes measured for leukaemia cells in peripheral blood.** The expression of four receptors was measured: FLT3, TIM3, CD33 and CD123. Scale bar: 5 μm.

**Table S3: Overview of the parameters derived from the DeepSPT analysis in this paper.**[1] * Is a metric added by the authors. For each parameter, except for density, the mean, median and standard deviation are taken into account.

| Name | Description | Unit |
|---|---|---|
| Alpha | Scaling exponent quantifying deviation from Brownian diffusion in the MSD–time relationship. Fitted per track on the MSD. | – |
| Diffusion coefficient (D) | The speed of diffusion found with the MSD fit. | $\mu m^2$ / s |
| P value | The P value of the MSD fitting, indicating the quality of the fit. | – |
| Efficiency | Quantifies path straightness by comparing net displacement to total path length. | – |
| logEfficiency | Logarithm of the Efficiency metric. | – |
| Fractal Dimensions | A mathematical value describing the capacity of a track to fill a space. | – |
| Gaussianity | Measure of how much the step length distribution deviates from a Gaussian. | – |
| Kurtosis | Kurtosis of the step length distribution, indicating the heaviness of the tails of the distribution. | – |
| MSD ratio | Approximate scaling exponent derived from comparing MSD values at two lag times. | – |
| Trappedness | The likelihood that a particle with a certain diffusion coefficient remains confined within an area over a period of time. | – |
| t0 | The percentage of steps spent in the slowest HMM state. | % |
| t1 | The percentage of steps spent in the second slowest HMM state. | % |
| t2 | The percentage of steps spent in the second fastest HMM state. | % |

| | | |
|---|---|---|
| lifetime | Mean number of steps spent in a HMM state. | - |
| avgSL | Average step length. | μm |
| avgMSD | Average MSD of a track. | $μm^2$ |
| AvgDP | Average dot product of successive displacement vectors, indicating directional persistence. | $μm^2$ |
| corrDP | Correlation of successive displacement directions. | – |
| signDP | Sign-based measure of whether steps tend to continue forward or reverse. | – |
| minSL | Minimum step length in the trajectory. | μm |
| maxSL | Maximum step length in the trajectory. | μm |
| BroadnessSL | Maximum step length minus minimum step length. | μm |
| covSL | Coefficient of variation of the step lengths (std/mean). | – |
| FractionSlow | Fraction of steps classified as slow. (Below 0.1 μm.) | % |
| Volume | The area or volume of the convex hull encompassing the localizations of trajectory. | $μm^2$ (2D) or $μm^3$ (3D) |
| perc_ND | Percentage of steps classified as normal diffusion. | % |
| perc_DM | Percentage of steps classified as directed motion. | % |
| perc_CD | Percentage of steps classified as confined diffusion. | % |
| perc_SD | Percentage of steps classified as subdiffusive motion. | % |
| num_changepoints | Count of transitions between predicted diffusion types. | - |
| Inst_D | Single step diffusion coefficient estimated with the formula MSD / 2dt. | $μm^2$ / s |
| meanSequence | Average value of the diffusion type labels predicted along the trajectory, where states are encoded as 0 (ND), 1 (DM), 2 (CD), and 3 (SD). | – |
| medianSequence | Median value of the sequence encoding. | – |
| maxSequence | Maximum value of the sequence encoding. | – |
| stdSequence | Standard deviation of the sequence encoding. | – |
| simSeq | Distance measure comparing diffusion type patterns, computed as the Euclidean distance between sequences encoded using 0 (ND), 1 (DM), 4 (CD), and 6 (SD). | – |
| confinement_ratio* | Net displacement divided by total path length; measures how straight the track is. | – |
| median_photon_count* | Median photon count of the track. | - |

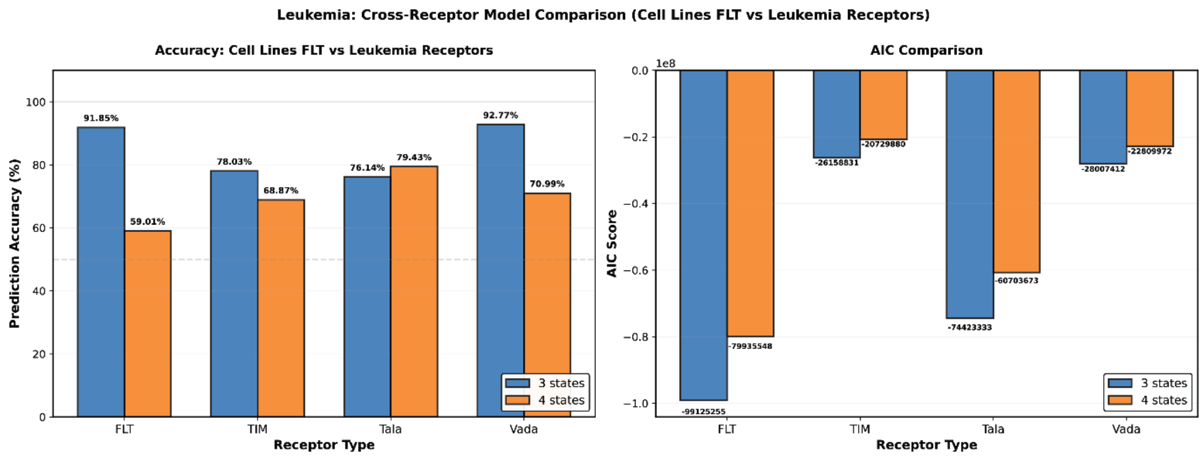


**Figure S5: Comparison of the 3-state and 4-state HMM model for all 4 receptors.** The accuracy of the 3 state model is higher than the 4 state model for FLT-3 and Vada, and the Akaike Information Criterion is lower than the 4-state model for all four receptors.

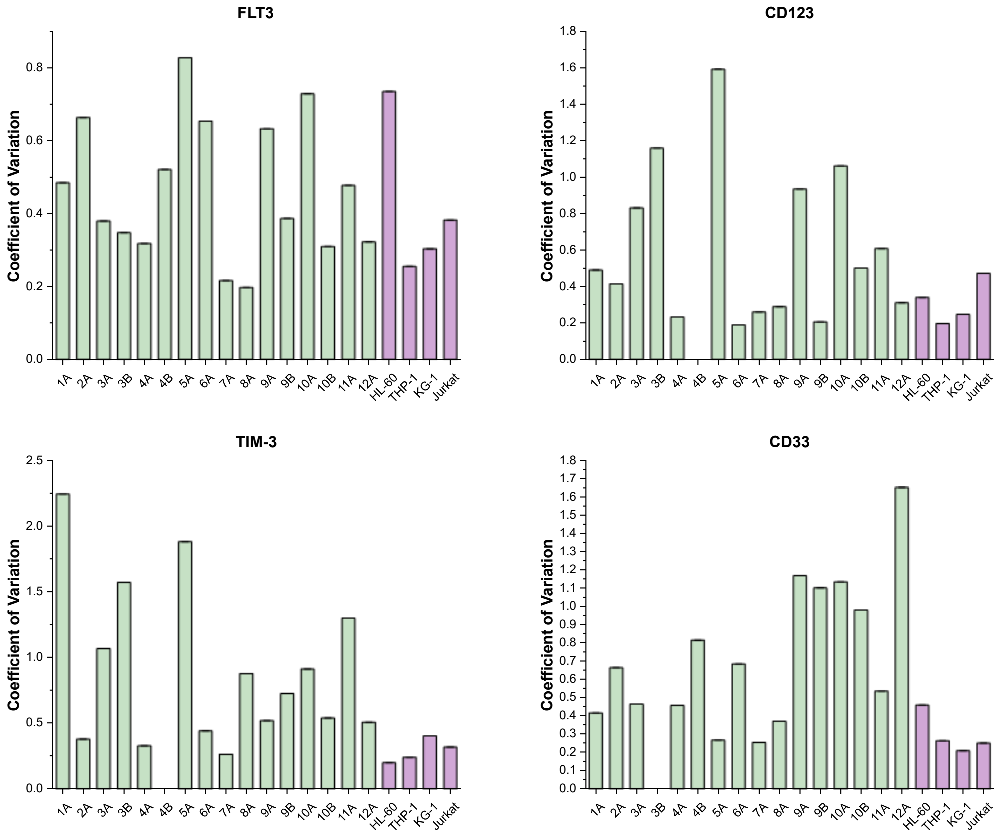


**Figure S6: Coefficients of variation of the densities per cell compared between patients and cell lines.** CD33 and TIM-3 are not statistically different (p=0.143 and 0.703, respectively), while FLT3 and CD123 are statistically different (p=0.014 and p= 0.037 respectively) when comparing patients vs cell lines.

### Extensive discussion about the relation between clinical data and receptor expression.

Interesting observations can be made for the different biomarkers and their correlation with the clinical data of the patients. FLT3 shows high density in several diagnostic samples, and certain subpopulations are enriched while others disappear during treatment. For patients 3 (samples 3A vs 3B) and 4 (4A vs 4B) this results in an overall decrease of FLT3 expression, while a significant increase is observed for patient 9. For patient 10, the average level of FLT3 expression remains the same, but the distribution becomes narrower, which is also reflected in the CV values. Receptor expression is currently not monitored over the course of treatment, which would be made possible with our approach and could potentially give insights in treatment resistance or changes in subpopulations. Usually, FLT3 expression is upregulated in AML and not in the myelodysplastic syndrome (MDS), which is supported by the low expression levels found for patient 6, who suffers from MDS.

TIM-3, by contrast, shows generally low surface abundance in the patients. CD123 expression varies considerably. This is in partial agreement with mutation status (e.g. higher levels in NPM1-mutant patients ). For the patients that were followed during their disease, patients 3B, 9B and 10B show increased CD123 compared to diagnosis. Notably, high CD123 in NPM1-mutant patients, such as patients 3 and 4, has been linked to improved outcomes when treated with venetoclax plus hypomethylating agents, providing potential therapeutic guidance. CD33 also displays variable expression across patients, but remains more stable over the course of treatment.

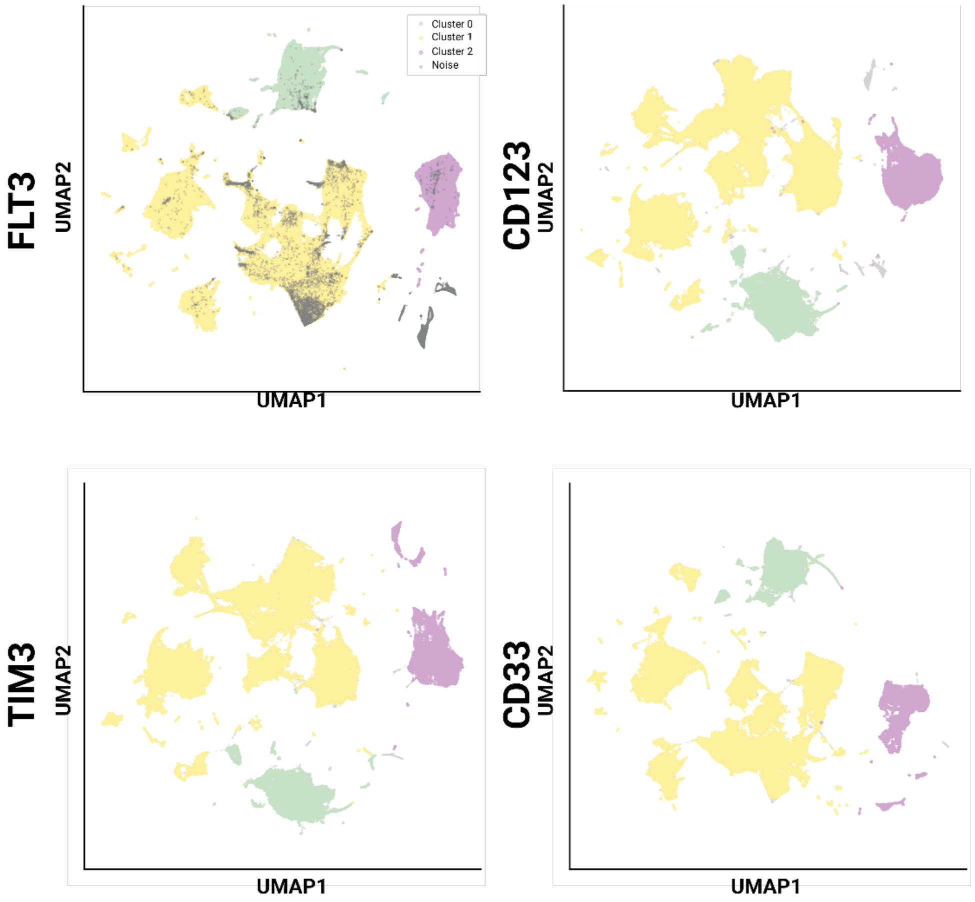


**Figure S7: Track level UMAPS for CD123, TIM3 and CD33 behaviour on all patients, a random 25% of the data was selected to plot. HDBSCAN was used for clustering the data.**

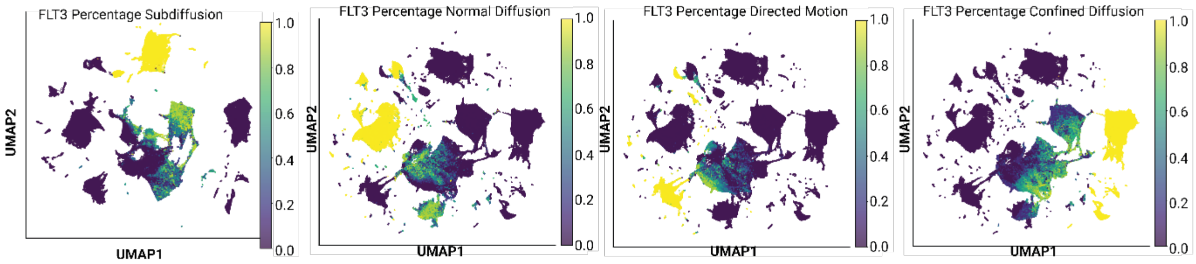


**Figure S8: Track level UMAPs of selected parameters clearly showing separation in the clusters.**

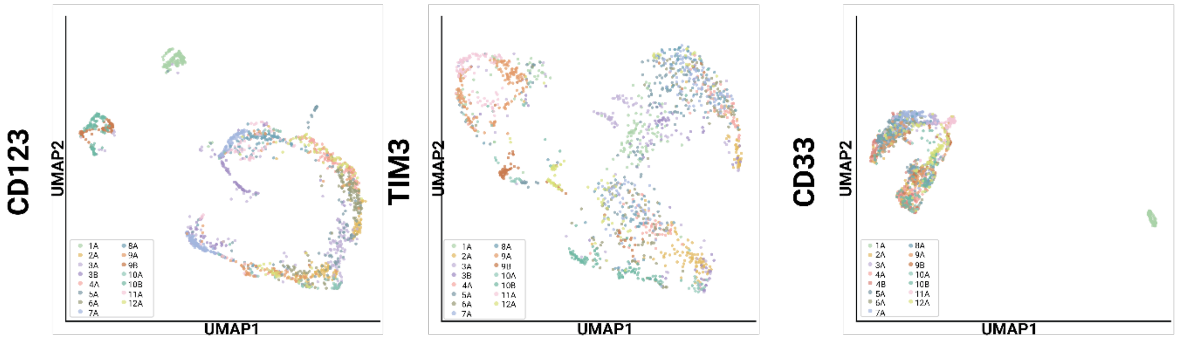


**Figure S9: Cell level UMAPS of the behaviour of CD123, TIM3 and CD33, colour coded by patient number.**

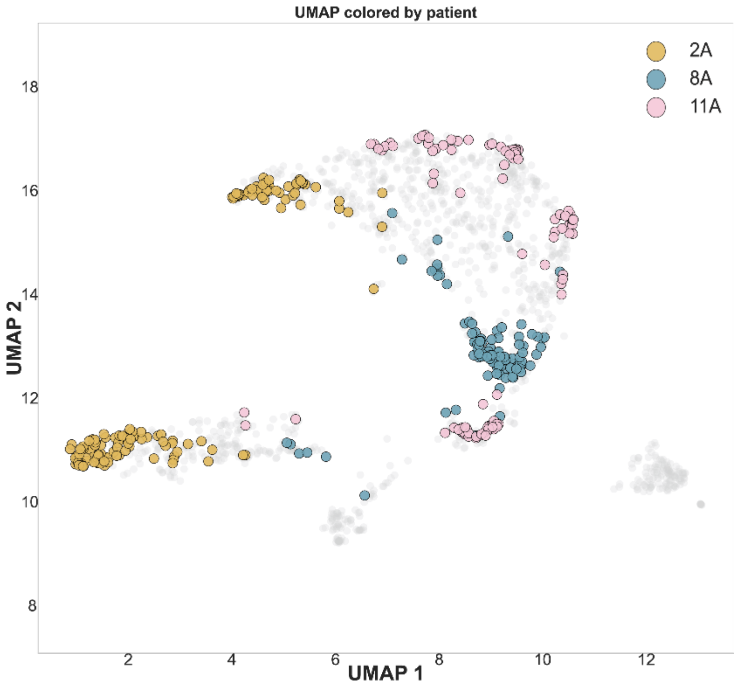


**Figure S10: UMAP of FLT3 with 3 selected patients highlighted, other patients are depicted in gray.** This demonstrates that individual patients are grouped together in the cluster and further away from other patients.

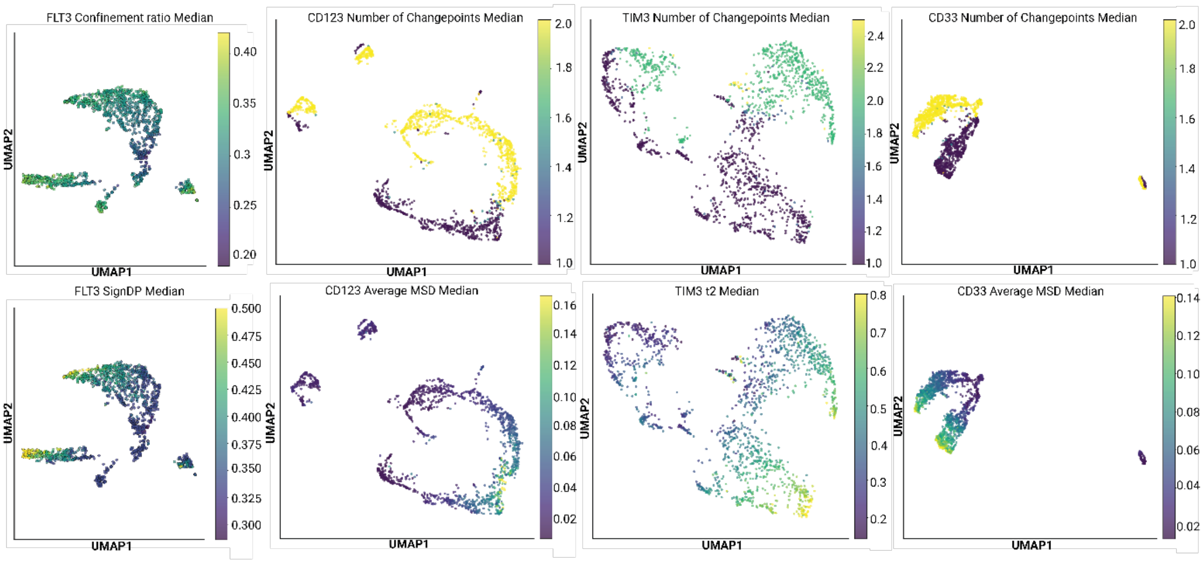


**Figure S11: Parameters indicating a spread in the UMAP for all four leukemia receptors and all patients, colour coded by the value of that parameter.**

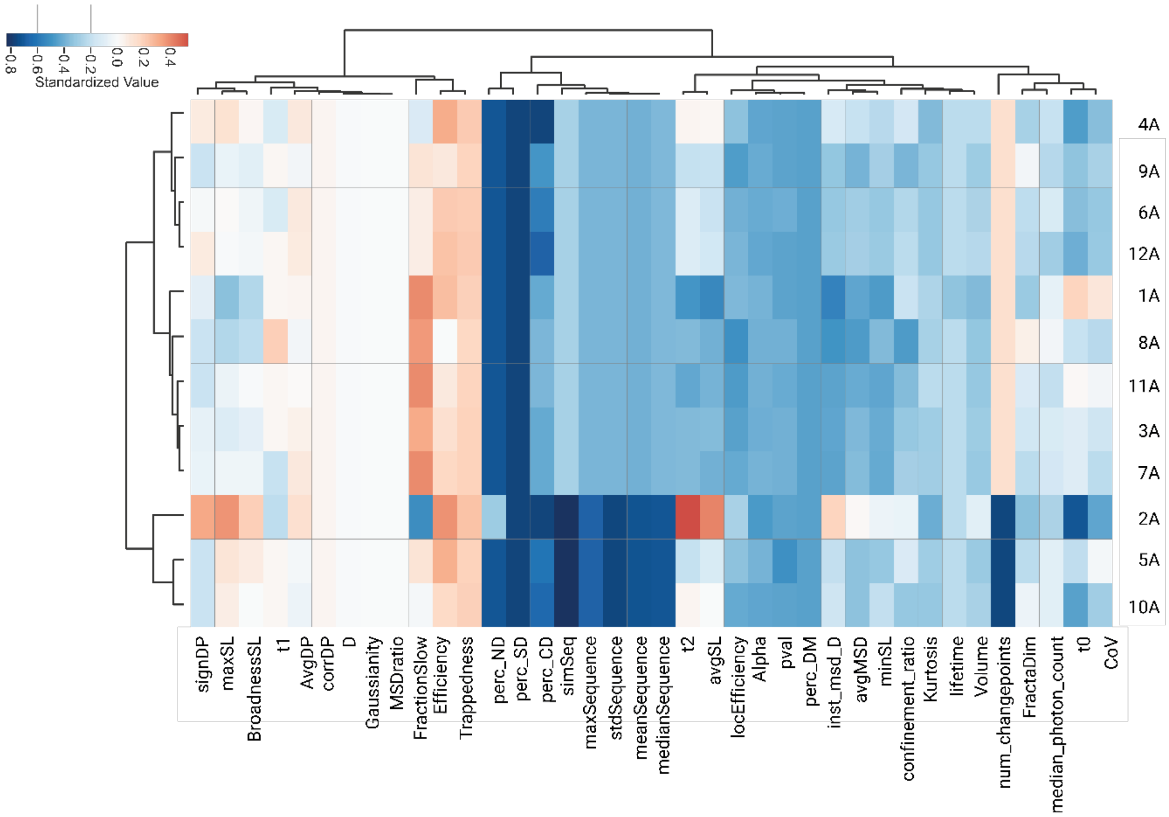


**Figure S12: Dendrogram of Fig. 5 with the parameters per column indicated.**

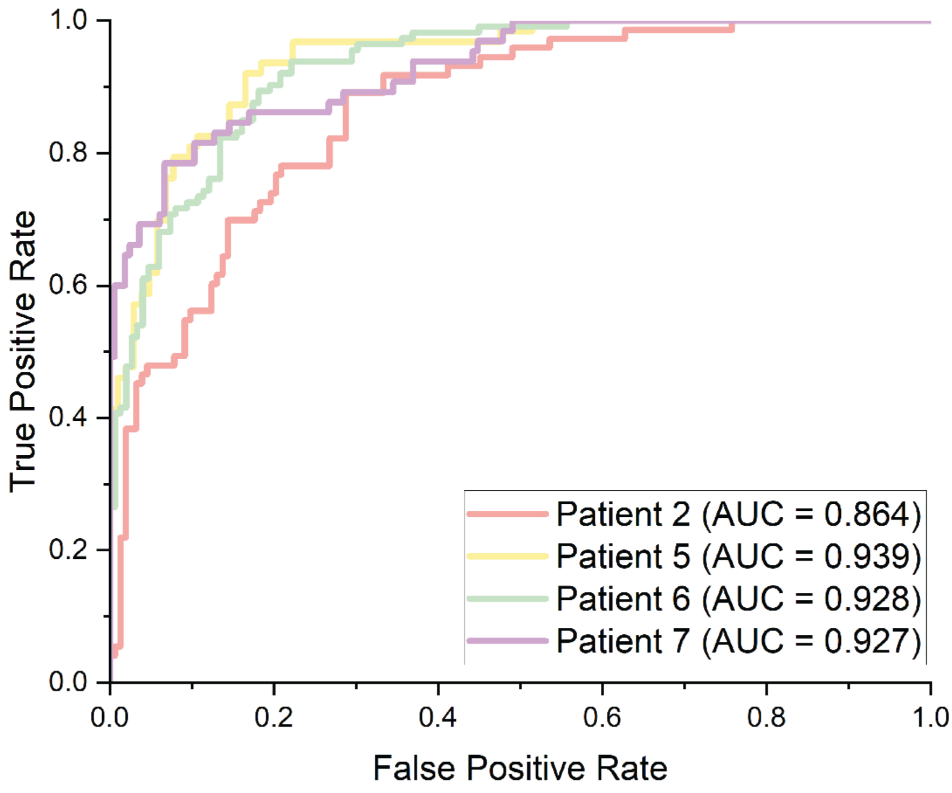


**Figure S13: XGBoost ROC Curves (by test patient)**

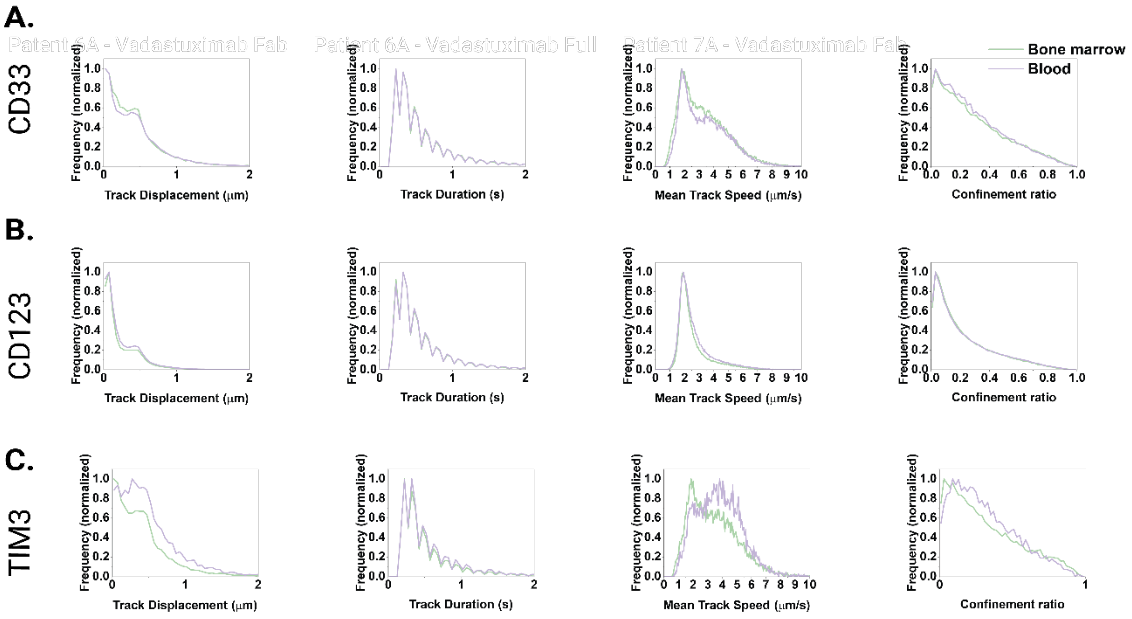


**Figure S14: Comparison of mobility parameters between peripheral blood and bone marrow blasts. A) CD33, B) CD123, C) TIM-3.**

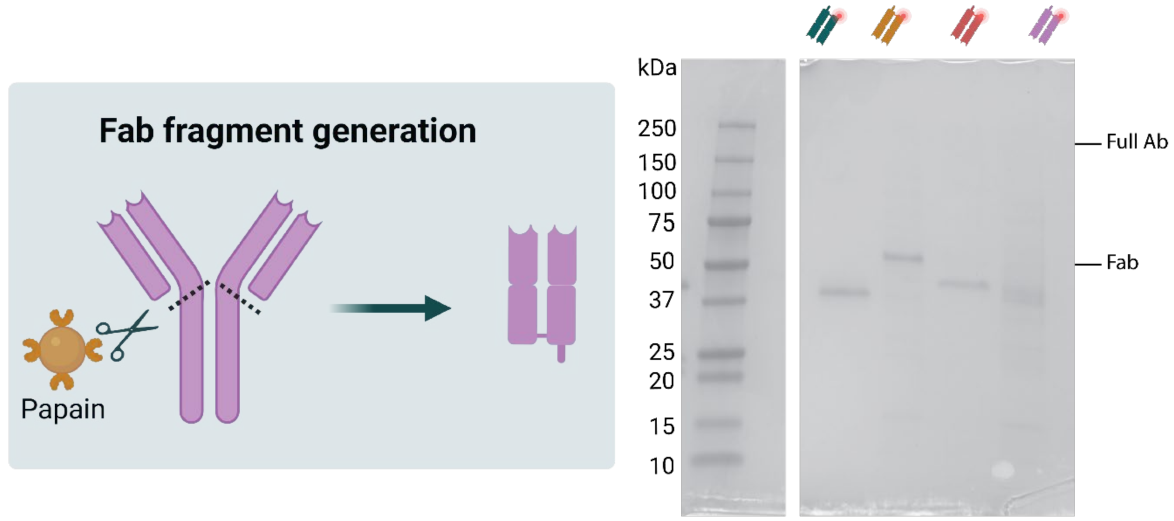


**Figure S15: Probe generation. The schematic shows papain digestion of antibodies. SDS-PAGE of all AML Fab fragments. Green Ab: anti-FLT3, orange: Talacotuzumab (CD123), red: anti-TIM-3, purple: Vadastuximab (CD33).**
